# Heightened subcortical reactivity to uncertain-threat is associated with future internalizing symptoms, conditional on stress exposure

**DOI:** 10.1101/2025.06.11.659182

**Authors:** Shannon E. Grogans, Kathryn A. DeYoung, Juyoen Hur, Allegra S. Anderson, Samiha Islam, Hyung Cho Kim, Jazmine Wedlock, Logan E. Craig, Rachael M. Tillman, Sanjana Das, Manuel Kuhn, Christopher C. Conway, Andrew F. Fox, Jason F. Smith, Alexander J. Shackman

**Affiliations:** Department of Psychology, University of Maryland, College Park, MD, USA; Neuroscience and Cognitive Science Program, University of Maryland, College Park, MD, USA; Maryland Neuroimaging Center, University of Maryland, College Park, MD, USA; Department of Psychology, Yonsei University, Seoul, Republic of Korea; Department of Psychiatry and Human Behavior, The Warren Alpert Medical School of Brown University, Providence, RI, USA; Department of Child and Adolescent Psychiatry and Behavioral Science, Children’s Hospital of Pennsylvania, Philadelphia, PA, USA; Manning College of Information and Computer Sciences, University of Massachusetts Amherst, Amherst, MA; River City Counseling, Chattanooga, TN, USA; McGill Neuropsychology, Bethesda, Maryland 20814 USA; Department of Psychology, University of Pittsburgh, Pittsburgh, PA, USA; Center for Depression, Anxiety and Stress Research, McLean Hospital, Harvard Medical School, Belmont, MA, USA; Department of Psychology, Fordham University, Bronx, NY, USA; Department of Psychology, Davis, CA, USA; California National Primate Research Center, Davis, CA, USA

**Keywords:** anxiety and depression, bed nucleus of the stria terminalis (BST/BNST), clinical affective neuroscience, extended amygdala, periaqueductal gray (PAG), prospective-longitudinal study design

## Abstract

**Background:** Anxiety, depression, and related internalizing illnesses are a leading burden on global public health, and often emerge during times of stress. Yet the underlying neurobiology has remained enigmatic, hindering treatment development.

**Methods:** Here we used a combination of tools—including a well-established threat-anticipation fMRI paradigm and longitudinal assessments of internalizing symptoms and negative life events (NLEs)—to identify the neural systems associated with future internalizing illness in a risk-enriched sample of 224 emerging adults followed for 2.5 years. We performed parallel analyses in an overlapping sample of 209 participants who completed a popular threat-related faces paradigm.

**Results:** Here we show that heightened reactivity to uncertain-threat anticipation in the bed nucleus of the stria terminalis and the periaqueductal gray is associated with a worsening longitudinal course of broadband internalizing symptoms among individuals with low levels of NLE exposure. These associations were specific to uncertain threat and generally remained significant when controlling for concurrent measures of threat-elicited distress or psychophysiological arousal, highlighting the added value of the neuroimaging measures. Symptom trajectories were unrelated to amygdala and frontocortical reactivity to anticipated threat. Contrary to past research, amygdala reactivity to threat-related faces was unrelated to future symptoms.

**Conclusions:** These observations provide a novel neurobiological framework for conceptualizing transdiagnostic internalizing risk and lay the groundwork for mechanistic and therapeutics research. A racially diverse, risk-enriched sample and pre-registered, best-practices approach enhance confidence in the robustness and translational relevance of these results.

## INTRODUCTION

Anxiety and depression lie on a continuum and, when extreme or pervasive, can become debilitating [1–6]. Internalizing-spectrum disorders are common—afflicting ∼700M individuals annually—and often co-occur [7–14]. Existing treatments are inconsistently effective [3, 5, 15–20]. Yet the neural systems that confer risk for internalizing illness remain elusive, impeding the development of improved interventions.

While the etiology of internalizing illness is undoubtedly complex and multifactorial [1, 21, 22], recent work highlights the importance of exaggerated reactivity to uncertain-threat anticipation. Heightened defensive responses to uncertain threat predict the onset and progression of internalizing symptoms, distinguish patients from controls, and are dampened by SSRIs and other treatments [23–37]. Converging lines of mechanistic and neuroimaging research have begun to reveal the neural systems governing responses to uncertain threat [38–42]. This work underscores the importance of subcortical regions, including the central nucleus of the amygdala (Ce), bed nucleus of the stria terminalis (BST), and periaqueductal gray (PAG). But it also highlights more recently evolved frontocortical regions, including the anterior insula (AI), frontal operculum (FrO), midcingulate cortex (MCC), and dorsolateral prefrontal cortex/frontal pole (dlPFC/FP). Cross-sectional research shows that heightened BST reactivity to uncertain threat is associated with individual differences in neuroticism/negative emotionality (N/NE)— a key dispositional risk factor for internalizing illness—but it cannot address whether BST reactivity is prospectively associated with future illness [43–47]. Meta-analytic research demonstrates that individuals with anxiety disorders show hyperreactivity to unpleasant emotional challenges in the amygdala, BST, PAG, MCC, and AI [41], but it remains possible that these neurobiological marks represent symptoms, scars, or concomitants of acute internalizing illness [48]. Addressing these fundamental questions mandates a prospective-longitudinal approach.

Negative life events (NLEs) precipitate, maintain, and exacerbate internalizing illness [49–58], but the brain circuits that serve as diatheses for stress-induced symptom progression remain poorly understood. In one of the only studies of its kind, Swartz, Hariri and colleagues showed that greater amygdala reactivity to ‘threat-related’ (fearful/angry) faces is associated with a worsening course of internalizing symptoms in undergraduates who self-reported greater NLE exposure. These findings highlight the potentially crucial role of experience-dependent pathways to psychopathology (e.g., *Amygdala × NLEs → Internalizing*). Whether these findings are reproducible, whether they extend to other threat-sensitive regions, and whether they generalize to genuinely distressing threats (e.g., shock) remains unknown.

Here we used a novel combination of tools—including a well-established certain/uncertain threat-anticipation fMRI paradigm and 30-month longitudinal assessments of internalizing symptoms and NLEs—to understand the neural circuits that promote internalizing illness in a racially diverse sample of 224 emerging adults, all of whom were free from internalizing illness at baseline (**Figure 1**). Participants were selectively recruited from a pool of 6,594 pre-screened individuals, ensuring a broad spectrum of dispositional risk and addressing a key limitation of convenience samples [59]. We focused on ‘emerging adulthood’ (∼18-22 years) because it is a time of profound, often stressful transitions [60–69]. More than half of undergraduate students report moderate-to-severe symptoms of anxiety and depression, with many experiencing the first emergence or recurrence of frank internalizing illness during this turbulent developmental chapter [10, 70–83]. This approach enabled us to test the hypothesis that increased recruitment of canonical threat-sensitive regions will prospectively predict the worsening of internalizing symptoms, that these associations will be more evident for uncertain than certain threat anticipation, and that these associations will be amplified by NLE exposure. To provide a direct link with prior work by Swartz and on-going biobank studies [84], we performed parallel analyses in an overlapping sample of 209 participants who completed a threat-related faces paradigm. Prospective associations with *DSM-5* internalizing diagnoses are described in the **Supplement.** Study hypotheses and general approach were pre-registered (https://osf.io/v6w79).

**Figure 1.**
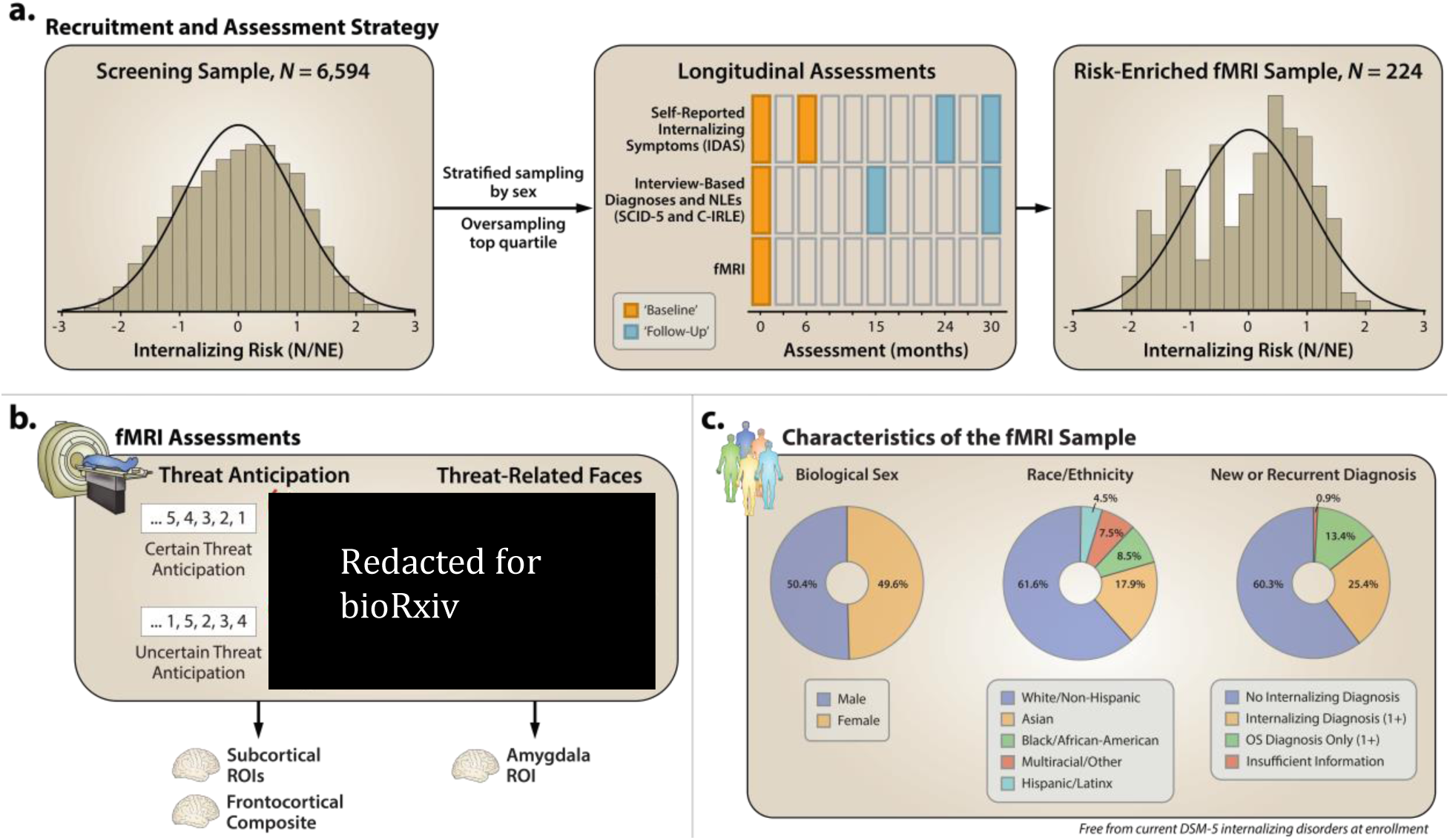
Study Overview. a. Recruitment and assessment strategy. To ensure a broad spectrum of N/NE, participants were selectively recruited from a racially diverse pool of 6,594 prescreened individuals. N/NE was assessed at initial screening and at the 0- and 6-month laboratory sessions. To maximize reliability and power, analyses leveraged a composite measure of N/NE that was aggregated across two scales and three measurement occasions. Top panels indicate the distribution (histogram) of N/NE in the screening (*left*) and risk-enriched fMRI samples (*right*). Middle panel indicates the intervals at which longitudinal assessments occurred. During the 30-month longitudinal follow-up, self-reported internalizing symptoms were assessed at 0, 6, 24, and 30 months using the IDAS. As in prior work by our group, the 0- and 6-month assessments were averaged to form an overall “baseline” composite and the 24- and 30-month assessments were averaged to form a “follow-up” composite. Internalizing diagnoses and NLEs were assessed at 0, 15, and 30 months using the SCID-5 and C-IRLE, respectively. fMRI was assessed at 0 months. **b. *Baseline fMRI assessments.*** *Threat anticipation.* All participants completed the Maryland Threat Countdown paradigm, a well-established anxiety–provocation paradigm. The paradigm takes the form of a 2 (Valence: Threat, Safety) × 2 (Temporal Certainty: Certain, Uncertain) repeated-measures design. On threat trials, subjects saw a stream of integers that terminated with the temporally certain or uncertain presentation of a noxious electric shock, unpleasant photograph, and thematically related audio clip. Safety trials were similar but terminated with the delivery of emotionally neutral stimuli. Hypothesis testing focused on activation associated with the anticipation of temporally certain and uncertain threat, relative to safety. A total of 220 individuals provided usable imaging data. *Threat-related faces.* An overlapping set of 213 participants also completed a “threat-related” (fearful/angry) faces fMRI paradigm. Participants viewed short blocks of photographs, alternating between blocks of faces and places (e.g., park, office). Hypothesis testing focused on activation associated with threat-related faces, relative to places. Consistent with prior work by our group, activation was quantified using subcortical and cortical ROIs and spatially unsmoothed fMRI data. **c. *Characteristics of the fMRI sample.*** Ring plots depict the relative frequencies of biological sex (*left*), race and ethnicity (*middle*), and the emergence of new or recurrent internalizing diagnoses among the combined fMRI sample of 224 participants (*right*). Abbreviations—C-IRLE, Cambridge Interview for Recent Life Events; *DSM-5,* Diagnostic and Statistical Manual of Mental Disorders, Fifth Edition; fMRI, functional magnetic resonance imaging; IDAS, Inventory of Depression and Anxiety Symptoms; N/NE, neuroticism/negative emotionality; NLEs, negative life events; SCID-5, Structured Clinical Interview for *DSM-5*.

## METHOD

### Overview

This project leverages data from the Maryland iRisk Study, a 30-month prospective-longitudinal study of a racially diverse sample of emerging adults (*n*=224; 49.55% female; 38.39% BIPOC). For full details, see the **Supplement**.

We used questionnaire measures of N/NE to screen 6,594 first-year university students [43, 85]. Screening data were stratified into quartiles, separately by sex. Given our focus, approximately half the participants were recruited from the top quartile, with the remainder split between the other quartiles (**Figure 1a**), enabling us to sample a broad spectrum of internalizing risk without discontinuities.

All participants self-reported adequate general health (see the **Supplement**) and were free from lifetime psychotic/bipolar disorders; current internalizing diagnosis; severe substance misuse; active suicidality; and on-going psychiatric treatment. Individuals with an adjustment disorder, ‘other specified’ internalizing disorders, or persistent depressive disorder who did not experience a major depressive episode in the past 2 months were not excluded. Participants provided informed written consent. Procedures were approved by the University of Maryland Institutional Review Board.

At baseline (0 months), the MTC (**Figures 1b, Supplementary Figure S1**) was used to quantify neural reactivity to the uncertain and certain anticipation of threat. Reactivity to a popular emotional-faces paradigm was also assessed (**Figures 1b, Supplementary Figure S3**). At 0, 6, 24, and 30 months, demographics and self-reported internalizing symptoms were assessed (**Figure 1a**). Diagnoses and NLEs were assessed using ‘gold-standard’ clinical interviews at 0, 15, and 30 months (**Figure 1a**).

### Participants

A total of 224 participants had usable data for one or both fMRI tasks (49.6% female; 61.6% White, 17.9% Asian, 8.5% African American, 4.5% Hispanic, 7.5% Multiracial/Other; *M*=18.8 years, *SD*=0.3; **Figure 1c**). See the **Supplement** for additional details.

### Internalizing symptoms

Internalizing symptoms were assessed using the Inventory of Depression and Anxiety Symptoms (IDAS) [86], modified to cover past-month symptoms. The IDAS Dysphoria scale, a 10-item broadband measure of transdiagnostic internalizing symptoms, served as the primary outcome [87–89]. We separately averaged the 0- and 6-month assessments to form a “baseline” composite and the 24- and 30-month assessments to form a “follow-up” composite (see the **Supplement**). Internal-consistency reliability was acceptable (α=0.91-0.93). Data missingness was minimal (**Supplementary Table S1**).

### NLEs

NLE exposure was assessed via semi-structured clinical interview, per best-practices [90]. Compared to questionnaires, interview measures reduce a number of common errors (e.g., false positives, false negatives, idiosyncratic event interpretations), resulting in more reliable and valid assessments of exposure frequency and severity [90, 91]. Here the Cambridge Interview for Recent Life Events (C-IRLE) was used to probe the occurrence and clinician-rated negative impact of 64 life events [92]. Negative impact was rated on a 1 (*severe*) to 4 (*mild*) scale. At each follow-up, NLEs were determined for the past 15 months. Consistent with past work [93], we computed an overall severity-weighted frequency score. Severity ratings were reverse scored—placing them on a 1 (*mild*) to 4 (*severe*) scale—and summed. Data missingness was minimal (**Supplementary Table S1**).

### Threat-anticipation paradigm

The MTC paradigm takes the form of a 2 (*Valence:* Threat, Safety) × 2 (*Temporal Certainty:* Uncertain, Certain) event-related design. For details, see the **Supplement**. On certain-threat trials, participants saw a descending stream of integers (“countdown”) for 18.75 s that culminated with the presentation of a noxious electric shock, unpleasant photograph, and aversive audio clip. Uncertain-threat trials were similar, but the integer stream was randomized and presented for an uncertain and variable duration (8.75-30.00 s; *M*=18.75 s). Safety trials were similar, but terminated with benign reinforcers. Prior work in this and other samples demonstrates that the MTC elicits robust symptoms of subjective distress and signs of objective arousal, and recruits the canonical threat-anticipation network, confirming its validity as an experimental probe of fear and anxiety [43, 94, 95].

### Threat-related faces paradigm

The threat-related faces paradigm takes the form of a pseudo-randomized block design. For details, see the **Supplement**. Participants viewed photographs of angry/fearful/happy faces, or emotionally neutral scenes. To ensure engagement, participants judged whether the current photograph matched the prior one.

### MRI data acquisition

MRI data were acquired using a Siemens Magnetom TIM Trio 3 Tesla scanner and standard techniques (see the **Supplement**). Sagittal T1-weighted images were acquired using a MPRAGE sequence (TR=2,400 ms; TE=2.01 ms; inversion=1,060 ms; flip=8°; thickness=0.8 mm; in-plane=0.8 × 0.8 mm; matrix=300 × 320; field-of-view=240 × 256). A co-planar T2-weighted image was also collected (TR=3,200 ms; TE=564 ms; flip=120°). A multi-band sequence was used to collect oblique-axial EPI volumes (acceleration=6; TR=1,250 ms; TE=39.4 ms; flip=36.4°; thickness=2.2 mm, slices=60; in-plane=2.1875 × 2.1875 mm; matrix=96 × 96). Two spin echo (SE) images were collected for fieldmap correction.

### MRI pipeline

Methods are identical to those described in prior work [43, 85, 95, 96]. Functional scans with excessive motion artifacts (>2 *SD*) were discarded. Participants with insufficient usable data (<2 scans of the threat-anticipation task or <1 scan of the threat-perception task) or who showed poor behavioral performance on the threat-perception task (accuracy <2 *SD*) were excluded from analyses.

#### First-level modeling

Modeling was identical to prior work [43]. For details, see the **Supplement**. Regressors were convolved with a canonical HRF. *Threat-anticipation.* Hemodynamic reactivity was modeled using variable-duration rectangular (“boxcar”) regressors that spanned the anticipation epochs of uncertain threat, certain threat, and uncertain safety. Certain-safety served as the reference condition and contributed to the baseline estimate [97]. *Threat-related faces.* Reactivity to blocks of each emotion was modeled using rectangular regressors. Place blocks served as the reference condition and contributed to the baseline estimate [97].

#### Second-level modeling

Whole-brain voxelwise repeated-measures GLMs were computed using *SPM12*. For threat-anticipation, we modelled activation during the anticipation of threat (vs. safety), uncertain threat (vs. baseline), and certain threat (vs. baseline). For threat-related faces, we modelled activation during the presentation of threat-related (angry and fearful) faces (vs. baseline).

#### Brain metrics

Activation was quantified using spatially unsmoothed data, maximizing resolution [43]. *Threat-anticipation, subcortical regions.* Ce, BST, and PAG activation were quantified using anatomical ROIs [98–100]. Regression coefficients were averaged across voxels and hemispheres for each combination of task contrast, region, and participant. *Threat-anticipation, frontocortical regions.* Frontocortical regions are large and functionally heterogeneous, rendering anatomical ROIs suboptimal. Here, frontocortical ROIs were prescribed based on peak task effects within anatomical regions previously identified in a large-scale neuroimaging study of threat anticipation [43], including MCC, AI, frontal operculum (FrO), and dorsolateral prefrontal cortex/frontal pole (dlPFC/FP). For each combination of region and hemisphere, regression coefficients were extracted using cubical ROIs (216 mm^3^) centered on the regional peak [85]. Regression coefficients were separately averaged for each combination of condition, region, and participant. To minimize the number of comparisons, we created composite measures of frontocortical activity for each condition (mean inter-region *r*=0.57; α>0.79). *Threat-related faces, amygdala.* The amygdala was anatomically defined. Regression coefficients were extracted for each hemisphere for the threat-related faces contrast and averaged for each participant.

### Analytic strategy

See the **Supplement** for full details. Regression models controlled for baseline Dysphoria. Separate models were implemented for each brain metric. Interactions were decomposed using simple-slopes [101]. The same approach was used to test whether associations are more evident for uncertain threat, controlling for certain threat. Associations with categorical internalizing diagnoses are described in the **Supplement**.

## RESULTS

### A substantial portion of the iRisk sample showed worsening of internalizing illness

Across the longitudinal follow-up, we observed marked differences in broadband internalizing symptoms. Nearly half the risk-enriched sample (45.1%) showed numeric worsening of dysphoria symptoms (**Figure 2a-b**) and more than a quarter (25.4%) developed a new or recurrent *DSM-5* internalizing diagnosis (**Figure 2c**). Together, these observations suggest that there is sufficient clinical change for meaningful brain-phenotype association analyses.

**Figure 2.**
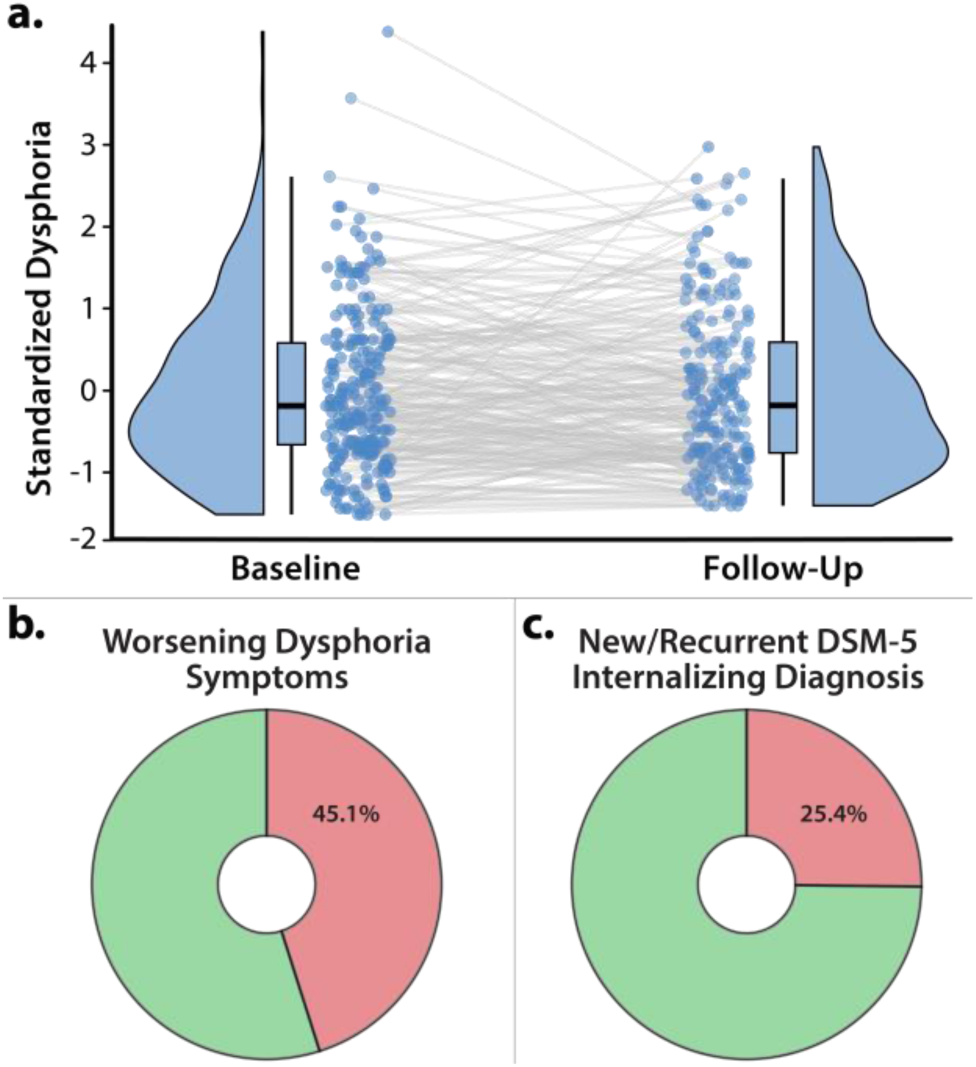
Longitudinal change in internalizing symptoms and diagnoses. **a. *Variation in internalizing symptom trajectories***. We observed marked individual differences in broadband internalizing symptoms. Nearly half (48.2%) of the sample showed at least a ±0.5 *SD* change between the baseline and follow-up composites (Baseline: *M*=20.28, *SD*=6.78; Follow-Up: *M*=20.93, *SD*=7.76; *r*(218)=0.63, *p*<0.001). Raincloud plots depict the smoothed density distributions (i.e., ‘bean’ or ‘half-violin’) of standardized Dysphoria. Box-and-whisker plots indicate the medians (*horizontal lines*) and interquartile ranges (*boxes*). Whiskers depict 1.5× the interquartile range. Colored dots connected by gray lines indicate changes in standardized N/NE for each participant. **b. *Worsening symptom trajectories.*** Nearly half of the sample (45.1%) showed numeric worsening of transdiagnostic Dysphoria symptoms across the 30-month longitudinal follow-up. **c. *Substantial emergence of new or recurrent internalizing diagnoses***. More than a quarter of the sample (25.4%) developed a new or recurrent *DSM-5* internalizing diagnosis. Abbreviation—DSM-5, Diagnostic and Statistical Manual, fifth edition.

### Marked differences in the quality and frequency of NLE exposure

Descriptively, our sample experienced NLEs across several key life domains, including work, relationships, and academics (**Figure 3a**). There was substantial variance in the frequency of NLE exposure (*M*=9.41, *SD*=3.66). Severity was largely confined to the mild-to-moderate range (*M*=1.14, *SD*=0.12). As expected, there was marked variation in the overall NLE composite used for hypothesis testing (*M*=10.77, *SD*=4.35; **Figure 3b**).

**Figure 3.**
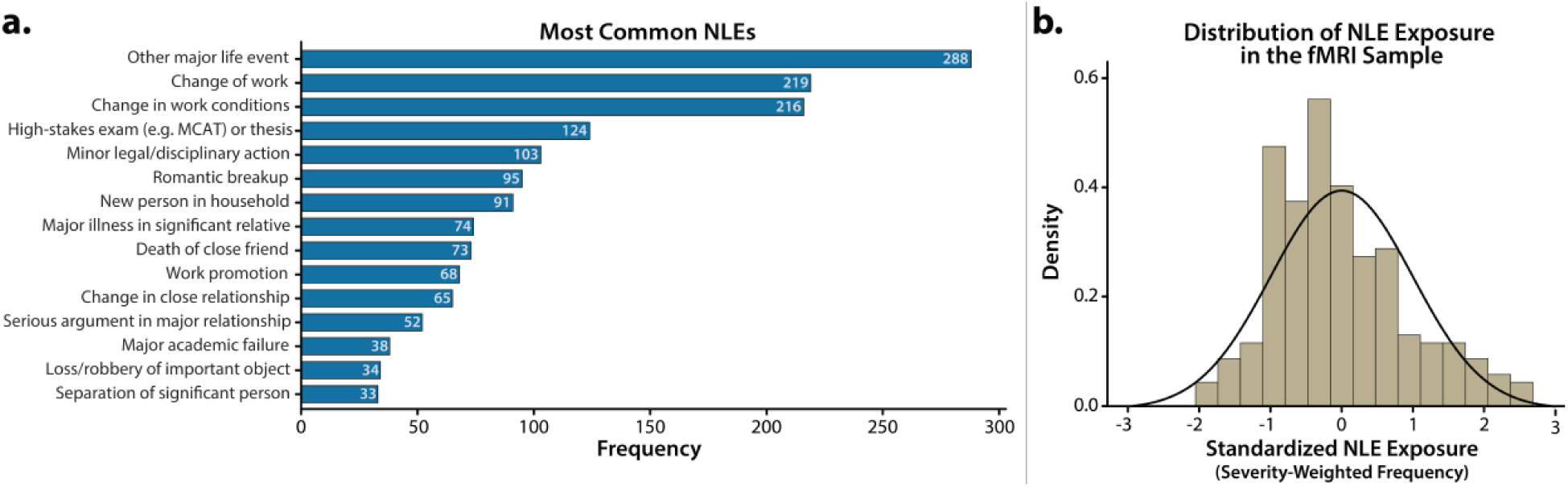
NLEs in the present sample. **a. *Most frequently reported NLEs.*** Bar chart depicts the 15 most frequently reported NLEs across the 30-month follow-up. **b. *Distribution of NLE exposure in fMRI sample.*** Histogram depicts the distribution of severity-weighted frequency of NLE exposure across the 30-month follow-up. Abbreviations—fMRI, functional magnetic resonance imaging; NLEs, negative life events.

### Both fMRI paradigms have the expected impact on brain function

We used one-sample *t*-tests to confirm that all ROIs (e.g., amygdala) evinced nominally significant activation for each of the relevant task contrasts, (*t*s(219)>3.08, *p*s<0.002; **Supplementary Table S2**).

### NLE exposure is associated with worsening internalizing symptoms

We used a simplified regression model (“base model”) to confirm that baseline Dysphoria and longitudinal NLE exposure are positively associated with Dysphoria at follow-up. Both prospective associations were significant (**Figure 4**). Elevated symptoms at baseline were associated with more severe Dysphoria at follow-up, controlling for NLEs (*β* =0.63, *t*(213)= 12.14, *p*<0.001, *R*^2^*_partial_*=0.409). Likewise, individuals exposed to more frequent or severe NLEs reported more severe Dysphoria at follow-up, controlling for baseline symptoms (*β*=0.20, *t*(213)=3.79, *p*<0.001, *R*^2^*_partial_*=0.063)—that is, they showed a worsening symptom trajectory across the 30-month follow-up. These observations underscore the potent consequences of NLEs for the longitudinal course of broadband internalizing symptoms and reinforce the general validity of our approach.

**Figure 4.**
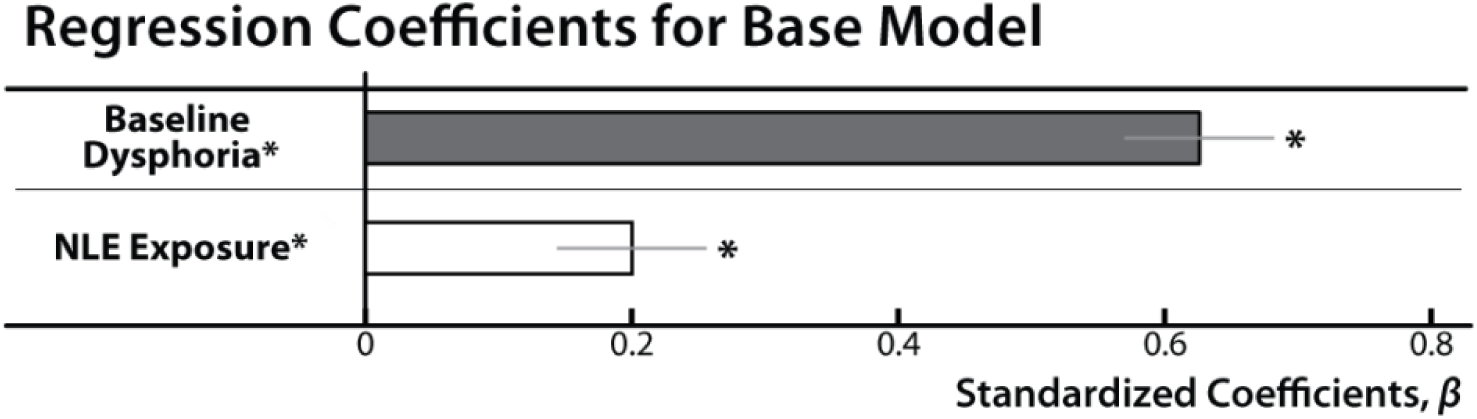
Summary of base linear regression models for IDAS Dysphoria analyses. Heightened broadband internalizing symptoms at baseline (*dark gray*) were associated with elevated symptoms at follow-up (*p*<0.001). Exposure to more severe or frequent NLEs (*white*) was also associated with greater internalizing symptoms at follow-up (*p*<0.001), consistent with prior work. Bars depict standardized coefficients for each regression model. Whiskers indicate standard errors. Significant associations are marked by an asterisk. Abbreviation—NLE, negative life event.

### Heightened BST and PAG reactivity to uncertain-threat anticipation is associated with worsening internalizing symptoms, conditional on NLE exposure

Building on our base model, we used a series of regressions to determine whether neural reactivity (e.g., BST) to anticipated threat—aggregating across certain and uncertain trials—is related to longitudinal changes in broadband internalizing symptoms. None of the associations were significant (*p*s>0.05; **Supplementary Table S3**).

Prior work suggests that the link between brain function and internalizing symptoms is likely to be stronger for measures of neural reactivity to uncertain-threat anticipation [46, 102]. To test this, we computed a series of regression models that included regional reactivity to uncertain and certain threat as simultaneous predictors. This provides an estimate of the variance in future internalizing symptoms uniquely explained by regional activation during each type of threat. Results revealed a significant interaction between BST reactivity to uncertain-threat and NLE exposure (*b*=-0.18, *t*(209)=-2.70, *p*=0.008, *R*^2^*_partial_*=0.034; **Figures 5-6 and Table 1**). The interaction remained significant when excluding BST_Certain-Threat_ as a predictor (*b*=-0.13, *t*(211)=-2.29, *p*=0.023, *R*^2^*_partial_*=0.024). For bivariate associations and additional descriptive statistics, see **Supplementary Tables S4-S5**. To decompose the significant interaction, we conducted a standard simple-slopes analysis. This revealed significant conditional effects of BST_Uncertain-Threat_ reactivity when NLE exposure was low (−1 *SD*: *b*=0.19, *t*(209)=2.23, *p*=0.027), but not when exposure was average (*b*=0.01, *t*(209)=0.17, *p*=0.86) or high (+1 *SD*: *b*=-0.17, *t*(209)=-1.79, *p*=0.07; **Figure 6**).

**Figure 5.**
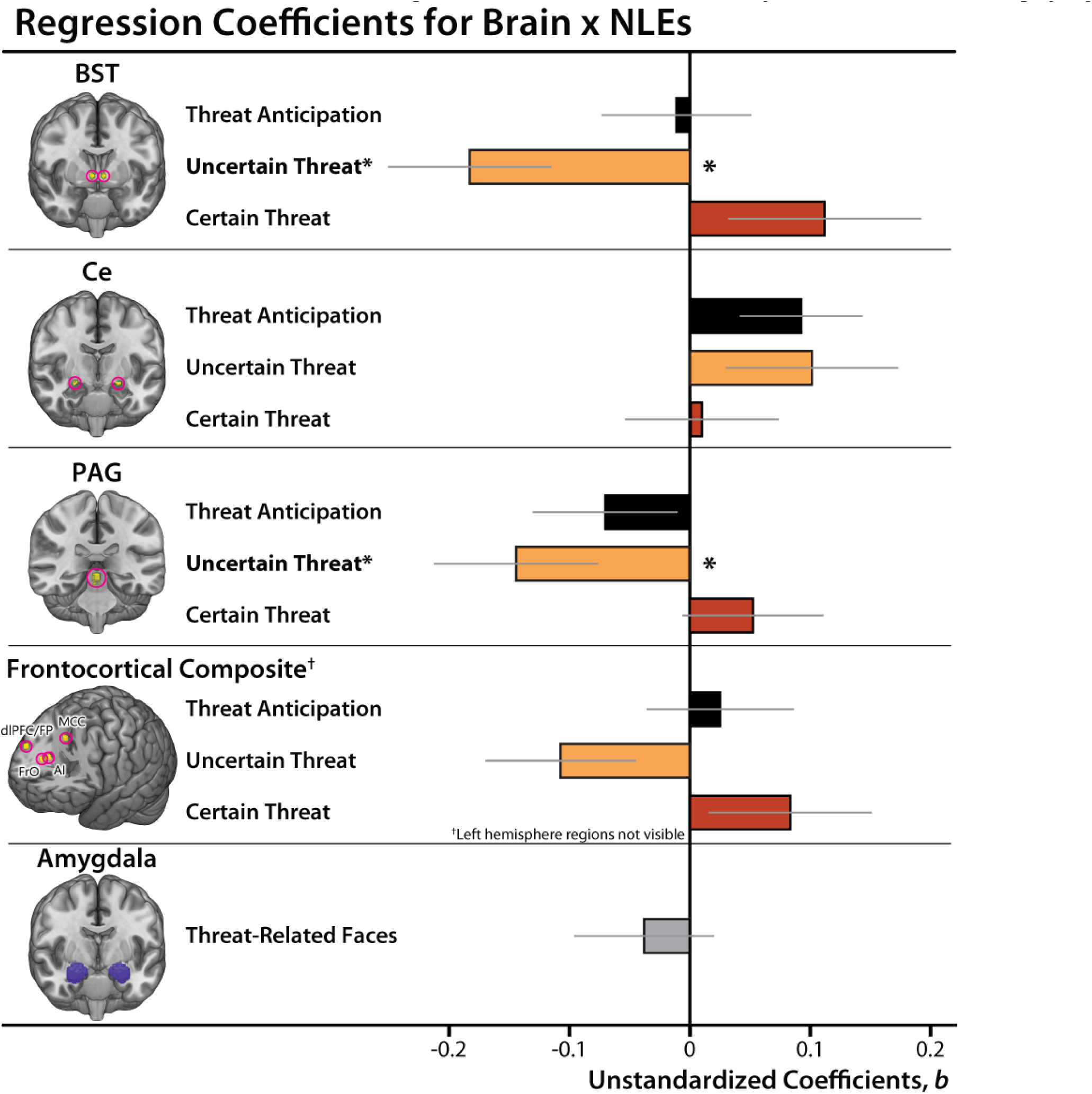
Summary of prospective Brain × NLE associations with Dysphoria. Bars depict unstandardized coefficients for each regression model. For detailed results, see **Table 4**. Whiskers indicate standard errors. Significant associations are indicted by asterisks. Abbreviations—AI, anterior insula; BST, bed nucleus of the stria terminalis; Ce, dorsal amygdala in the region of the central nucleus of the amygdala; dlPFC/FP, dorsolateral prefrontal cortex/frontal pole; FrO, frontal operculum; MCC, midcingulate cortex; PAG, periaqueductal gray; NLEs, negative life events.

**Figure 6.**
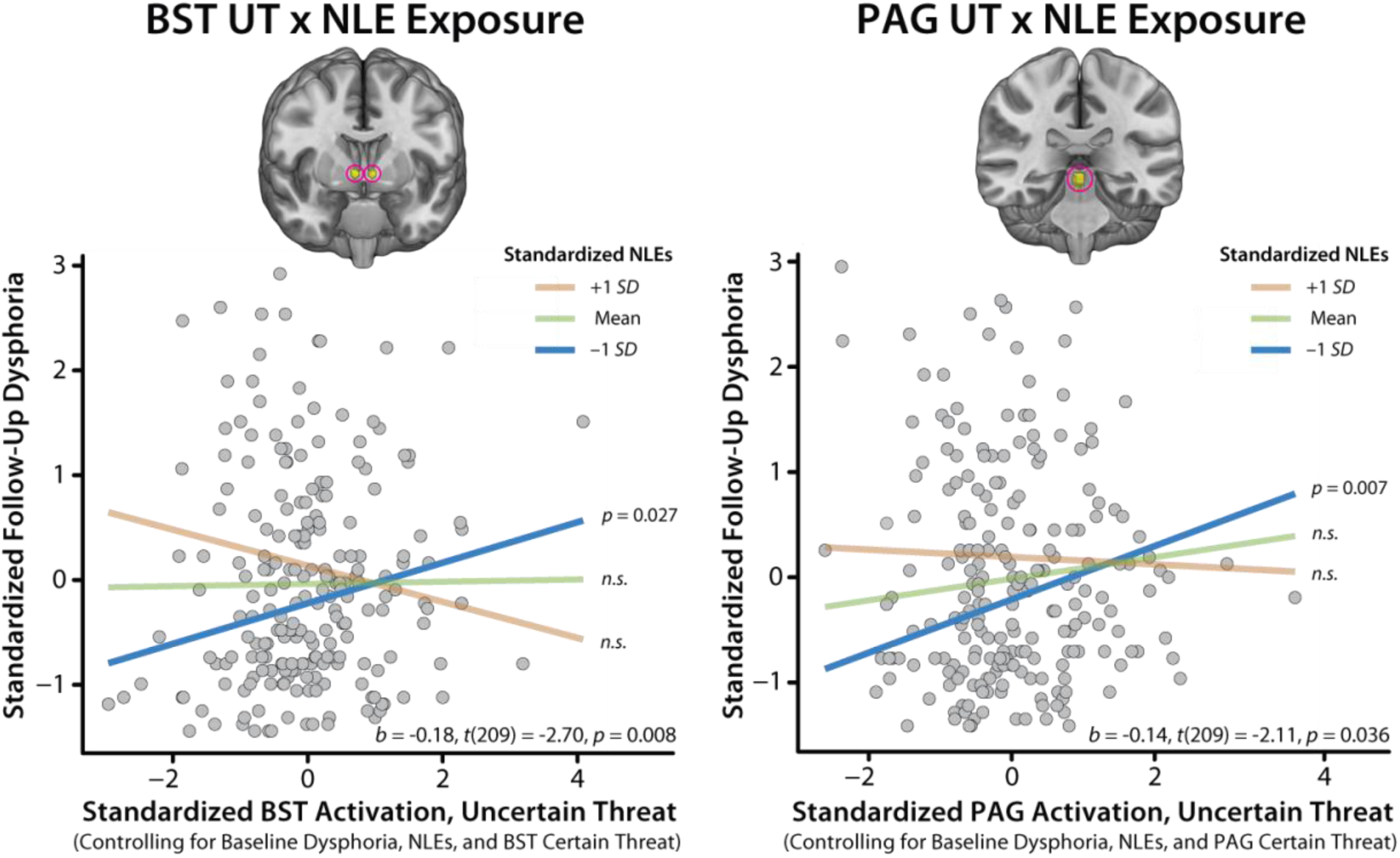
Simple-slopes analyses for the BST/PAG_Uncertain-Threat_-NLE interactions. Greater BST/PAG_Uncertain-Threat_ reactivity was associated with significant longitudinal worsening of broadband Dysphoria symptoms among individuals with low (−1 *SD*; *blue*) NLE exposure, but not among those with average (*green*) or high (+1 *SD*; *orange*) exposure. This figure is for illustrative purposes only; hypothesis tests leveraged continuous measures and were computed using the entire sample. Abbreviations— BST, bed nucleus of the stria terminalis; NLEs, negative life events; PAG, periaqueductal gray.

**Table 1.** Linear regressions of follow-up Dysphoria on neural reactivity to the anticipation of threat or the presentation of threat-related faces, NLEs, and their interaction, controlling for baseline Dysphoria.

| Model Terms <sup>a,b</sup> | *b* | *t* | *df* | *p* | *R*<sup>2</sup><sub>partial</sub> |
| --- | --- | --- | --- | --- | --- |
| **BST** | — | — | — | — | — |
| NLEs *** | 0.18 | 3.53 | 209 | <0.001 | 0.056 |
| Baseline Dysphoria *** | 0.63 | 11.87 | 209 | <0.001 | 0.403 |
| BST, Uncertain-Threat Anticipation<sup>c</sup> | 0.01 | 0.17 | 209 | 0.863 | 0.000 |
| BST, Certain-Threat Anticipation<sup>c</sup> | -0.02 | -0.36 | 209 | 0.716 | 0.001 |
| BST UT<sup>c</sup> × NLEs ** | -0.18 | -2.70 | 209 | 0.008 | 0.034 |
| BST CT<sup>c</sup> × NLEs | 0.11 | 1.40 | 209 | 0.164 | 0.009 |
| **Ce** | — | — | — | — | — |
| NLEs *** | 0.20 | 3.86 | 209 | <0.001 | 0.067 |
| Baseline Dysphoria *** | 0.63 | 12.14 | 209 | <0.001 | 0.413 |
| Ce, Uncertain-Threat Anticipation<sup>c</sup> | 0.03 | 0.38 | 209 | 0.702 | 0.001 |
| Ce, Certain-Threat Anticipation<sup>c</sup> | -0.07 | -0.98 | 209 | 0.327 | 0.005 |
| Ce UT<sup>c</sup> × NLEs | 0.10 | 1.41 | 209 | 0.159 | 0.009 |
| Ce CT<sup>c</sup> × NLEs | 0.01 | 0.15 | 209 | 0.884 | 0.000 |
| **PAG** | — | — | — | — | — |
| NLEs *** | 0.20 | 3.92 | 209 | <0.001 | 0.068 |
| Baseline Dysphoria *** | 0.63 | 12.15 | 209 | <0.001 | 0.414 |
| PAG, Uncertain-Threat Anticipation<sup>c</sup> | 0.11 | 1.72 | 209 | 0.087 | 0.014 |
| PAG, Certain-Threat Anticipation<sup>c</sup> | -0.11 | -1.71 | 209 | 0.089 | 0.014 |
| PAG UT<sup>c</sup> × NLEs * | -0.14 | -2.11 | 209 | 0.036 | 0.021 |
| PAG CT<sup>c</sup> × NLEs | 0.05 | 0.90 | 209 | 0.371 | 0.004 |
| **Frontocortical Composite** | — | — | — | — | — |
| NLEs *** | 0.19 | 3.65 | 209 | <0.001 | 0.060 |
| Baseline Dysphoria *** | 0.63 | 11.95 | 209 | <0.001 | 0.406 |
| Frontocortical Composite, UT<sup>c</sup> | 0.03 | 0.51 | 209 | 0.613 | 0.001 |
| Frontocortical Composite, CT<sup>c</sup> | -0.04 | -0.58 | 209 | 0.561 | 0.002 |
| Frontocortical Composite UT<sup>c</sup> × NLEs | -0.11 | -1.72 | 209 | 0.087 | 0.014 |
| Frontocortical Composite CT<sup>c</sup> × NLEs | 0.08 | 1.23 | 209 | 0.220 | 0.007 |
| **Amygdala** | — | — | — | — | — |
| NLEs *** | 0.18 | 3.38 | 204 | <0.001 | 0.053 |
| Baseline Dysphoria *** | 0.63 | 11.93 | 204 | <0.001 | 0.411 |
| Amygdala, Threat-Related Faces | -0.06 | -1.15 | 204 | 0.249 | 0.006 |
| Amygdala, Threat-Related Faces × NLEs | -0.04 | -0.65 | 204 | 0.516 | 0.002 |

<sup>a</sup>Model intercept terms were not significant ( $p > 0.05$ ). <sup>b</sup>All predictors were standardized (z-transformed). To sidestep standardizing the data twice, unstandardized coefficients are reported. <sup>c</sup>Relative to the implicit baseline (i.e., certain-safety). Abbreviations—BST, bed nucleus of the stria terminalis; Ce, dorsal amygdala in the region of the central nucleus; CT, Certain-Threat anticipation; PAG, periaqueductal gray; ROI, region of interest; UT, Uncertain-Threat anticipation. \* $p \leq 0.05$ , \*\* $p < 0.01$ , \*\*\* $p < 0.001$ .

The same general pattern was evident for the PAG (*PAG_Uncertain-Threat_ × NLE: b*=-0.14, *t*(209)=-2.11, *p*=0.036, *R*^2^*_partial_*=0.021; **Figures 5-6 and Table 1**). Here, the interaction was no longer significant when excluding PAG_Certain-Threat_ as a predictor (*b*=-0.11, *t*(211)=-1.90, *p*=0.059, *R*^2^*_partial_*=0.017). For bivariate associations and additional descriptive statistics, see **Supplementary Tables S6-S7**. Mirroring the BST, simple-slopes analyses revealed significant conditional effects of PAG_Uncertain-Threat_ reactivity when NLE exposure was low (−1 *SD*: *b*=0.25, *t*(209)=2.73, *p*=0.007), but not when exposure was average (*b*=0.11, *t*(209)=1.72, *p*=0.087) or high (+1 *SD*: *b*=-0.04, *t*(209)=-0.37, *p*=0.71; **Figure 6**). There were no significant effects for any of the other regions considered (**Figure 5**, **Table 1**, and **Supplementary Tables S8-S13**). Collectively, these observations underscore the explanatory value of BST and PAG reactivity. These associations were specific to uncertain threat and generally remained significant when controlling for concurrent measures of threat-elicited distress or psychophysiological arousal, highlighting the added value (“incremental validity”) of the neuroimaging measures (see the **Supplement**).

### Amygdala reactivity to threat-related faces is unrelated to future internalizing symptoms

We conducted parallel analyses for amygdala reactivity to threat-related faces. In contrast to prior work [93], amygdala reactivity was unrelated to the longitudinal course of internalizing symptoms, regardless of NLE exposure (*Amygdala × NLEs*: *p*=0.52; *Amygdala*: *p*=0.25; **Figure 5**, **Table 1, and Supplementary Tables S12-S13**). To clarify interpretation of the non-significant interaction, we computed standardized Bayesian effect sizes [103]. From a Bayesian perspective, the amygdala-NLE interaction showed weak evidence of an association with future internalizing symptoms (*BF_10_*=1.23). In contrast, the BST_Uncertain-Threat_-NLE interaction showed very strong evidence (*BF_10_*=38.20) and the PAG_Uncertain-Threat_-NLE interaction showed moderate evidence (*BF_10_*=9.17). Consistent with this overall pattern, the BST/PAG_Uncertain-Threat_-NLE associations both remained significant when controlling for amygdala reactivity to threat-related faces and the amygdala-NLE interaction (*BST × NLEs*: *b*=-0.19, *t*(196)=-2.75, *p*=0.007; *PAG × NLEs*: *β*=-0.14, *t*(196)=-2.10, *p*=0.037).

## DISCUSSION

Internalizing illnesses afflict hundreds of millions annually, and often emerge in response to stressors. Yet the underlying neurobiology has remained enigmatic, hindering treatment development. Here we show that heightened BST/PAG reactivity to uncertain-threat anticipation is associated with a worsening course of internalizing symptoms across the two-year follow-up (*BF_10_*=9.17-38.20). Contrary to expectation, these associations selectively manifested among individuals with relatively low NLE exposure (**Figure 6**). They were specific to uncertain threat and generally remained significant when controlling for concurrent measures of threat-elicited distress or psychophysiological arousal, highlighting the added value of the neuroimaging measures. Symptom trajectories were unrelated to Ce and frontocortical reactivity to anticipated threat. Contrary to past research, amygdala reactivity to threat-related faces was unrelated to future symptoms (*BF_10_*=1.23).

These results provide the first prospective-longitudinal evidence that heightened BST/PAG reactivity to uncertain threat foretells future internalizing symptoms, providing an important extension of prior cross-sectional research. That work shows that BST reactivity to uncertain threat is associated with elevated dispositional risk for internalizing illness (high N/NE) but it could not address whether variation in BST reactivity is associated with future internalizing illness [43]. Our findings show that it is. Conversely, a recent meta-analysis showed that the BST and PAG are hyper-reactive to unpleasant emotional challenges in individuals with anxiety disorders [41], but it could not rule out the possibility that these neurobiological marks represent scars or concomitants of acute illness [48]. Our findings, which leveraged neuroimaging data from individuals who were free from internalizing disorders at baseline, weigh against this possibility.

Our findings are broadly consistent with BST/PAG neurobiology. Both regions are recruited in neuroimaging studies of threat-anticipation [41, 94, 95]. Metabolism in both regions is associated with variation in dispositional anxiety in monkeys [104]. Both are anatomically poised to integrate diverse sources of potentially threat-relevant information and orchestrate specific signs of anxiety and other emotions [105–108]. A key challenge for the future will be to clarify the causal contribution of these regions to the development of internalizing illness. Rodent models show that the BST exerts bidirectional control over defensive responses to uncertain threat [109–113]. Traditionally, the PAG has been regarded as a simple downstream relay governed by the extended amygdala [105, 106, 108]. Yet emerging evidence suggests that the PAG can independently control defensive responses [108, 114]. Consistent with this perspective, micro-stimulation of the human PAG has been shown to evoke intense feelings of fear, elevated arousal, and a marked reluctance to continue stimulation [108, 115]. Moving forward, it will be useful to determine whether existing treatments dampen BST/PAG reactivity to uncertain threat and whether this depends on treatment modality (i.e., psychosocial vs. pharmacological). Research integrating fMRI with smartphone experience-sampling has the potential to clarify the momentary emotional processes that link heightened BST/PAG reactivity to internalizing illness [85].

Our findings demonstrate that exaggerated BST/PAG reactivity to uncertain threat is associated with a worsening trajectory of internalizing symptoms. Yet this prospective association was only evident among individuals with low levels of NLE exposure, saturating at higher levels. What does this imply? Anxiety and depression are complex and heterogeneous, both in presentation and etiology [116–119]. The present results suggest that the neural circuits underlying the development of broadband internalizing symptoms can be fractionated into two divisions, with some regions (BST/PAG) conferring biological risk in low-risk settings, and other—yet to be discovered regions—transducing higher levels of life stress into illness. Put another way, heightened BST/PAG to uncertain threat reactivity is only maladaptive when it occurs inpsychologically benign (“inappropriate”) environments [120, 121]. Similar dose-dependent liabilities have been documented by Kendler, Akil, and others in the psychiatric genetics literature (*Gene* × *Low-Adversity* → *Internalizing*) [68, 122]. This kind of dose-dependent biological “hand-off” also has clear conceptual precedents in the neuroscience literature. This work demonstrates that different “doses” of threat trigger qualitatively distinct defensive responses (e.g., vigilance vs. flight) [123]. This specificity is thought to reflect the selective recruitment of distinct neural circuits at different points along the threat intensity continuum [123, 124].

We and others have shown that internalizing symptoms and diagnoses are less common in low-stress environments (e.g., **Figure 4**). Their occurrence in these benign contexts may reflect a more “endogenous” etiology [117, 118]. Consistent with this perspective, BST/PAG reactivity to uncertain threat is associated with the heritable variation in dispositional anxiety (“nature”) in monkeys and with trait-like variation in neuroticism in humans [43, 106]. An important avenue for future research will be to determine whether elevated BST/PAG reactivity is associated with a more recurrent, endogenous lifetime course of internalizing illness.

We did not replicate a landmark report showing that heightened amygdala reactivity to threat-related faces is associated with a worsening course of broadband internalizing symptoms (∼15-22 months) among Duke University students with above-average levels of self-reported NLE exposure (*R^2^_partial_*=0.03) [93]. The authors framed the amygdala as a “neural biomarker of psychological vulnerability to future life stress.” We did not replicate these associations. Methodological differences between the two studies were modest and, if anything, seem to underscore the enhanced rigor of our approach (e.g., <3% attrition over 30 months, interview-based NLE assessments, cutting-edge fMRI pipeline). Importantly, we demonstrated substantial longitudinal change in internalizing illness (**Figure 2**) and robust amygdala activation to faces (**Supplementary Table S2**). Power to detect long-term symptom trajectories was similar across studies (Maryland iRisk, 30 months: *n*=209-218; Duke Neurogenetics, ∼22 months: *n*=192). On balance, these considerations cast doubt on claims that the amygdala represents a robust biomarker for future internalizing symptoms. They suggest that this prospective association is either narrower than originally envisioned, conditional on subtle differences in methodology, or non-existent. More broadly, the present results add to growing evidence that amygdala reactivity to threat-related faces is negligibly associated with internalizing risk or diagnostic status [43, 125].

Clearly, several challenges remain for the future. First, it will be important to determine whether our conclusions generalize to more representative samples and other types of threat. It will also be useful to explore associations with narrower internalizing symptoms [44, 126, 127]. Second, future work may benefit from more intensive NLE sampling and from disaggregating stressor type, frequency, and severity. Third, the effects of stress, adversity, and trauma are multidimensional, nonlinear, interactive, and age-dependent [128]. Unraveling this complexity is important, but it will require large, deeply phenotyped longitudinal cohorts. Fourth, animal models will be critical for generating hypotheses about the molecules, cell types, and microcircuits supporting variation in BST/PAG function [38, 39, 108, 129]. Fifth, the etiology of internalizing psychopathology is complex, multifactorial, and likely reflects the coordinated interactions of distributed neural networks [21, 22, 118, 130]. Moving forward, it will be important to determine the relevance of functional connectomics [131, 132]. For additional considerations, see the **Supplement**.

Internalizing illness imposes a staggering burden on public health. In the U.S., nearly 1 in 3 individuals will experience a lifetime anxiety disorder [133] and 1 in 5 will experience a depressive disorder [7]. Diagnoses and service utilization are surging [134–142]. Existing psychological and pharmacological treatments remain inaccessible, unacceptable (e.g., stigma), or intolerable for many [5, 18, 143–151], underscoring the urgency of developing a deeper understanding of the underlying neurobiology. Our findings demonstrate that heightened BST/PAG reactivity to uncertain-threat anticipation is associated with a worsening trajectory of broadband internalizing symptoms among individuals with low levels of NLE exposure. In contrast, amygdala reactivity was unrelated to future symptoms. These observations provide a neurobiological framework for conceptualizing transdiagnostic internalizing risk and lay the groundwork for mechanistic and therapeutics research. A comparatively large, racially diverse, and risk-enriched sample and a pre-registered, best-practices approach enhance confidence in the robustness and translational relevance of these results.

## ACKNOWLEDGMENTS

Authors acknowledge assistance and critical feedback from three anonymous reviewers, A. Antonacci, J. Blanchard, L. Friedman, J. Furcolo, M. Gamer, A. Gard, C. Grubb, G. Hancock, R. Hum, C. Kaplan, T. Kashdan, J. Kuang, C. Lejuez, D. Limon, B. Nacewicz, L. Pessoa, S. Rose, J. Swayambunathan, A. Vogel, members of the Affective and Translational Neuroscience laboratory, the staff of the Maryland Neuroimaging Center, and the Office of the Registrar at the University of Maryland. This work was partially supported by the California National Primate Center; National Institutes of Health (AA030042, AA031261, DA040717, MH107444, MH121409, MH121735, MH128336, MH129851, OD011107, MH131264, MH126426); National Research Foundation of Korea (2021R1F1A1063385 and 2021S1A5A2A03070229); University of California, Davis; University of Maryland; and Yonsei Signature Research Cluster Program (2021-22-0005). Authors declare no conflicts of interest.

## AUTHOR CONTRIBUTIONS

A.J.S., K.A.D., and J.F.S. designed the overall study. S.E.G. and A.J.S. envisioned the present project. J.F.S. and A.J.S. developed and optimized the imaging paradigm. K.A.D. managed data collection and study administration. K.A.D., J.F.S., A.S.A, S.I., J.W., L.E.C., and R.M.T. collected data. J.F.S. and M.K. developed data processing and analytic software for imaging analyses. J.H., J.F.S., and H.C.K. processed imaging data. S.E.G. and S.J. cleaned clinical diagnosis and life stress data. S.E.G., J.F.S., and A.J.S. analyzed data. S.E.G., C.C.C., and A.J.S. developed the analytic strategy. S.E.G., A.S.F., J.F.S., and A.J.S. interpreted data. S.E.G. and A.J.S. wrote the paper. S.E.G. created figures and tables. A.J.S. funded and supervised all aspects of the study. All authors contributed to reviewing and revising the paper and approved the final version.

## PREREGISTRATION

Our general approach and hypotheses were preregistered (https://osf.io/v6w79).

## RESOURCE SHARING

Raw data and select materials are publicly available at the National Institute of Mental Health Data Archive (https://nda.nih.gov/edit_collection.html?id=2447). Key neuroimaging maps are available at NeuroVault (https://neurovault.org/collections/13109/). De-identified processed data and analytic code are available via the Open Science Framework (https://osf.io/ufs7j/files/osfstorage).

