## Supplement for "Heightened subcortical reactivity to uncertain-threat is associated with future internalizing symptoms, conditional on stress exposure"

**Overview, recruitment, and general procedures**

The present project leverages archival data from the Maryland iRisk Study (R01-MH107444). iRisk is a 30-month prospective-longitudinal study of a racially diverse sample of emerging adults (*n*=224; 49.55% female; 38.39% BIPOC). Given the low level of data missingness (<2.68%; see **Supplementary Table S1**), specialized missing-data procedures (e.g., multiple imputation, full-information maximum likelihood estimation) were not employed.

The general study design was inspired by Alloy and Abramson's (1999) seminal 30-month Temple–Wisconsin study of depression in university students [1] and reflected a compromise between the scientific goal of tracking the participants for as long as possible—to enable greater opportunity for meaningful change in the severity of internalizing symptoms—and practical considerations, including the need to screen, enroll, and perform multiple waves of follow-up assessments within the constraints of a 5-year grant and 4-year baccalaureate degree program (**Figure 1a** in the main report).

We used well-established questionnaire measures of N/NE [2-4]—a key risk factor for the development of internalizing disorders—to screen 6,594 first-year university students (57.1% female; 59.0% White, 19.0% Asian, 9.9% African American, 6.3% Hispanic, 5.8% Multiracial/Other; *M*=19.2 years, *SD*=1.1 years) [5, 6]. Screening data were stratified into quartiles (top quartile, middle quartiles, bottom quartile), separately for males and females. Individuals who met preliminary inclusion criteria were independently and randomly recruited via email from each of the resulting six strata. Because of our focus on psychiatric risk, approximately half the participants were recruited from the top quartile, with the remainder split between the middle and bottom quartiles (i.e., 50% high, 25% medium, and 25% low; **Figure 1a** in the main report). This enabled us to sample a broad spectrum of internalizing risk without gaps or discontinuities, while balancing the inclusion of men and women. Simulation work suggests that this oversampling ('enrichment') approach does not bias statistical tests to a degree that would compromise their validity [7].

All participants had normal or corrected-to-normal color vision, and reported the absence of lifetime neurological symptoms, pervasive developmental disorder, very premature birth, medical conditions that would contraindicate MRI, and prior experience with noxious electrical stimulation. All participants were free from a lifetime history of psychotic and bipolar disorders; a current diagnosis of an anxiety, depressive, or trauma disorder (past 2 months); severe substance misuse; active suicidality; and on-going psychiatric treatment as determined by an experienced, masters-level diagnostician using the Structured Clinical Interview for *DSM-5* [SCID-5; 8]. To maximize the range of risk, subjects with a lifetime history of internalizing disorders were *not* excluded, nor were individuals who met criteria for adjustment disorder or 'other specified' internalizing disorders [1, 9]. Along these same lines, individuals who met criteria for persistent depressive disorder who did not experience a major depressive episode in the past 2 months were not excluded. Participants provided informed written consent and all procedures were approved by the Institutional Review Board at the University of Maryland, College Park (Protocol #659385).

At baseline (0 months), the Maryland Threat Countdown fMRI paradigm (**Figures 1b** in the main report, **Supplementary Figure S1**) was used to quantify neural reactivity to the uncertain and certain anticipation of threat (i.e., aversive stimulation). To provide a more direct link with on-going biobank research, reactivity to a popular emotional-faces paradigm was also assessed (**Figures 1b** in the main report, **Supplementary Figure S3**). At 0, 6, 24, and 30 months, self-reported internalizing symptoms were assessed (**Figure 1a** in the main report). Diagnoses and NLEs were assessed using 'gold-standard' clinical interviews at 0, 15, and 30 months, allowing us to test hypothesized moderators (e.g., NLEs) during a period of peak risk for some of the most common and debilitating internalizing illnesses (**Figure 1a** in the main report). Follow-up assessments were conducted in the laboratory or online according to participant preference.

Data from this study have been featured in prior work focused on relations between personality and the longitudinal course of internalizing symptoms [10], the basic neurobiology of fear and anxiety [11-13], relations between social anxiety and real-world mood [14], relations between threat-related brain activity and real-world mood dynamics [6], the neuroanatomical correlates of early-life anxiety shyness and behavioral inhibition [15], and relations between threat-related brain activity and N/NE [5], but have never been used to address the present aims.

**Participants**

A total of 241 participants were recruited and scanned. Of these, 6 withdrew due to excess distress in the scanner, 1 withdrew from the study after the imaging session, and 4 were excluded due to incidental neurological findings. A total of 224 participants had usable data for either one or both fMRI tasks, as outlined in more detail below (49.6% female; 61.6% White, 17.9% Asian, 8.5% African American, 4.5% Hispanic, 7.5% Multiracial/Other; *M*=18.8 years, *SD*=0.3; **Figure 1c** in the main report). Data missingness for the questionnaire and interview assessments is also described below.

***Threat-anticipation task.*** One participant was excluded from fMRI analyses due to gross susceptibility artifacts in the echoplanar imaging (EPI) data, 2 were excluded due to insufficient usable data (see below), 6 were excluded due to excess motion artifact (see below), and 1 was excluded due to task timing issues, yielding a racially diverse final sample of 220 participants (49.5% female; 61.4% White, 18.2% Asian, 8.6% African American, 4.1% Hispanic, 7.3% Multiracial/Other; *M*=18.8 years, *SD*= 0.4 years). Of these, 2 participants were excluded from skin conductance analyses due to insufficient usable data (see below). A subset of 209 participants (95.0%) provided usable data for both fMRI tasks.

***Threat-related faces task.*** Three participants were excluded due to gross susceptibility artifacts in the EPI data, 1 was excluded due to insufficient usable data (see below), 7 were excluded due to excessive motion artifact (see below), and 6 participants for inadequate behavioral performance (see below), yielding a final sample of 213 participants (49.3% female; 61.0% White, 17.8% Asian, 8.5% African American, 4.2% Hispanic, 7.0% Multiracial/Other; *M*=18.8 years, *SD*=0.3 years). A subset of 209 participants (98.1%) provided usable data for both fMRI tasks.

#### Power analyses

Sample size was determined *a priori* as part of the award that supported data collection (R01-MH107444) using simple benchmark (i.e., analysis independent) effect sizes. The target sample size ($N$≈240) was chosen to afford acceptable power and precision given available resources. At the time of study design, G-power (version 3.1.9.2) indicated >99% power to detect a benchmark ('generic') medium-sized effect ($r$=0.30) with up to 20% planned attrition ($n$=192 usable datasets) using α=0.05, two-tailed [16]. In practice, the final longitudinal sample of 216 usable multi-measure datasets (baseline threat-anticipation fMRI data and follow-up NLE and internalizing symptom data) provides 75% power to detect focal associations as small as $r$=0.177 ($R^2$=0.031).

#### Pre-registration and resource sharing

The general approach and predictions were pre-registered (https://osf.io/v6w79). Raw data and select materials are publicly available at the National Institute of Mental Health Data Archive (https://nda.nih.gov/edit_collection.html?id=2447). Key neuroimaging maps are available at NeuroVault (https://neurovault.org/collections/13109/). De-identified processed data and analytic code are available via the Open Science Framework (https://osf.io/ufs7j/files/osfstorage).

**Questionnaire assessments**

***Demographics.*** At baseline, participants self-reported age, race and ethnicity, and biological sex.

***Symptoms.*** *Assessment.* Internalizing symptoms were assessed using the Inventory of Depression and Anxiety Symptoms [17]. The IDAS includes 10 specific symptom scales: Appetite Gain, Appetite Loss, Ill Temper, Insomnia, Lassitude, Panic, Social Anxiety, Suicidality, Traumatic Intrusions, and Well-Being), as well a broadband measure of internalizing symptoms, Dysphoria [17, 18]. Variation in Dysphoria is a sensitive and specific marker of *DSM-5* internalizing diagnoses [19] and this scale served as the primary IDAS outcome variable. Participants used a 1 (*not at all*) to 5 (*extremely*) scale to rate themselves on a total of 64 items. Item responses were summed for each symptom scale. For the present study, the IDAS timeframe was modified to cover past-month symptoms. Participants completed the IDAS at 0, 6, 24, and 30 months. *Data reduction.* We created composite internalizing symptom measures by aggregating across adjacent pairs of longitudinal assessments. Specifically, we separately averaged the 0- and 6-month assessments to form a "baseline" composite and the 24- and 30-month assessments to form a "follow-up" composite. The decision to focus on these composites was motivated by a combination of conceptual and methodological considerations. Conceptually, we sought to understand the prospective relevance of threat-related brain function to changes in internalizing symptoms across the transition from late adolescence to early adulthood—a transition that spans years, not weeks or months [20]. Shorter-term fluctuations in internalizing symptoms were not central to the present aims. Given this goal, it was methodologically appealing to aggregate the two natural pairs of assessments—baseline (0 and 6 months) and follow-up (24 and 30 months)—with an eye to enhancing reliability and power [21-27]. For each pair of assessments, 6-month test-retest reliability was acceptable ($ICC_{3,1}$=0.74-0.79). Internal-consistency reliability was acceptable for both the baseline and follow-up composite ($\alpha$=0.91-0.93).

**Interview assessments**

***General protocol.*** Clinical interviews were conducted by a highly experienced masters-level diagnostician at 0, 15, and 30 months.

***Internalizing diagnoses.*** *Assessment.* Internalizing diagnoses were assessed using the SCID-5 [8]. Skip-out rules, which would prevent a diagnostic section or module from being fully administered, were omitted. At baseline, lifetime and current internalizing diagnoses were determined. At each follow-up assessment, internalizing diagnoses were determined for the prior 15 months. *Data reduction.* A binary Any Internalizing Diagnosis variable (*present/absent*) was created to capture the onset or recurrence of one or more internalizing illnesses during the longitudinal follow-up, including acute stress disorder, adjustment disorder, agoraphobia, generalized anxiety disorder, major depressive disorder, panic disorder, persistent depressive disorder, post-traumatic stress disorder, social anxiety disorder, and specific phobia. Adjustment disorder was included if: **(a)** full diagnostic criteria were not met at baseline, but were met at a follow-up interview, or **(b)** full criteria were met at baseline, remission was evident at the 15-month interview, and full criteria were again met at the 30-month interview. Likewise, persistent depressive disorder was included if full criteria were not met at the baseline clinical interview, but were met at a follow-up interview. Other-specified, substance-induced, medication-induced, and general-medical diagnoses were excluded.

***NLEs.*** *Assessment.* NLE exposure was assessed via semi-structured clinical interview, consistent with methodological best-practices [28]. Compared to self-report NLE questionnaires, interview-based measures reduce a number of common errors (e.g., false positives, false negatives, idiosyncratic event interpretations), resulting in a more reliable and valid assessment of exposure frequency and severity [28, 29]. Here the Cambridge Interview for Recent Life Events (C-IRLE) was used to probe the occurrence and clinician-rated negative impact of 64 distinct life events (e.g., *death of close friend, major academic failure, romantic breakup*) [30]. Negative impact was rated on a 1 (*severe*) to 4 (*mild*) scale, where lower ratings indicate greater severity. Here, negative impact was defined as "the degree of unpleasant impact, stress or threat the event would be expected to bring to bear on someone when its full nature and particular circumstances are taken into account," and clinicians were instructed not to let the individual's subjective report of the impact of an event unduly influence their rating [30]. At baseline, NLEs were reviewed for the prior two months. At the follow-up assessments, NLEs were determined for the past 15 months. *Data reduction.* Consistent with past work [31], we computed an overall severity-weighted frequency score. To do so, severity ratings were reverse scored—placing them on a 1 (*mild*) to 4 (*severe*) scale—and summed. Higher values indicate more frequent and/or severe NLE exposure during the 30-month follow-up.

**Questionnaire and interview data missingness and exclusions**

Data missingness for the questionnaire (IDAS) and interview assessments (SCID-5 and C-IRLE) was minimal (**Supplementary Table S1**).

**Threat-anticipation paradigm**

***Paradigm structure and design considerations.*** The Maryland Threat Countdown paradigm is a well-established, fMRI-optimized variant of temporally uncertain-threat assays that have been validated using fear-potentiated startle and acute anxiolytic administration (e.g., benzodiazepine) in mice, rats, and humans [32-36]. The paradigm has been successfully used in several prior fMRI studies [5, 6, 12, 37]. As shown in **Supplementary Figure S1**, the MTC paradigm takes the form of a 2 (*Valence:* Threat, Safety) × 2 (*Temporal Certainty:* Uncertain, Certain) randomized, event-related, repeated-measures design (3 scans; 6 trials/condition/scan). Subjects were completely informed about the task design and contingencies prior to scanning. Simulations were used to optimize the detection and deconvolution of task-related hemodynamic signals. Stimulus presentation and ratings acquisition were controlled using Presentation software (version 19.0, Neurobehavioral Systems, Berkeley, CA).

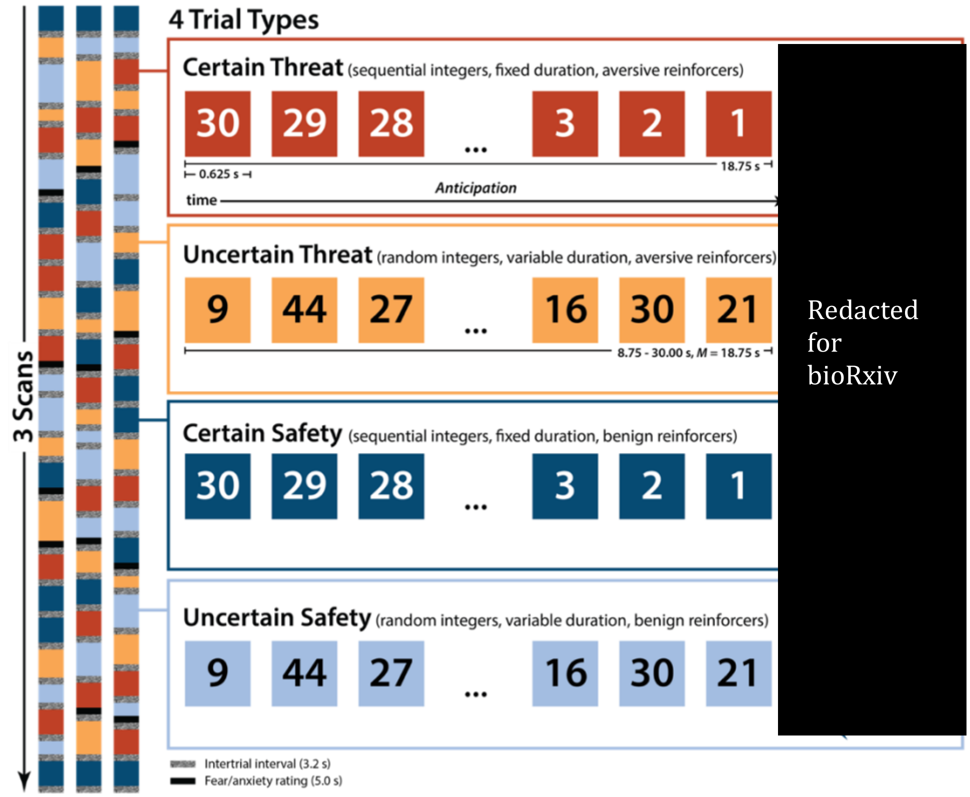

**Supplementary Figure S1. *Threat-anticipation paradigm.*** The Maryland Threat Countdown task takes the form of a 2 (*Valence:* Threat, Safety) × 2 (*Temporal Certainty:* Certain, Uncertain) repeated-measures, randomized event-related design. Participants were completely informed about the task design and contingencies prior to scanning. The task was administered in 3 scans, with short breaks between scans. On certain-threat trials, participants saw a descending stream of integers ("count-down") for 18.75 s. To ensure robust anxiety, this anticipation epoch always terminated with the presentation of a noxious electric shock, unpleasant photograph, and thematically related audio clip (e.g., scream). Uncertain-threat trials were similar, but the integer stream was randomized and presented for an uncertain and variable duration (8.75-30.00 s; *M*=18.75 s). Participants knew that something aversive was going to occur, but they had no way of knowing precisely *when*. Safety trials were similar, but terminated with the delivery of benign reinforcers (e.g., just-perceptible electrical stimulation). White-noise visual masks (3.2 s) were presented between trials to minimize persistence of visual reinforcers in iconic memory. Participants were periodically prompted to rate the intensity of fear/anxiety experienced a few seconds earlier, during the anticipation ('countdown') epoch of the prior trial. Skin conductance was continuously acquired.

On certain threat trials, participants saw a descending stream of integers ("count-down;" e.g., 30, 29, 28...3, 2, 1) for 18.75 s. To ensure robust distress, this anticipation epoch culminated with the presentation of a noxious electric shock, unpleasant photograph (e.g., mutilated body), and thematically related audio clip (e.g., scream, gunshot). Uncertain threat trials were similar, but the integer stream was randomized and presented for an uncertain and variable duration (8.75-30.00 s; *M*=18.75 s). Participants knew that something aversive was going to occur, but they had no way of knowing precisely when. Consistent with recent recommendations [38], the average duration of the anticipation epoch was identical across conditions, ensuring an equal number of measurements (TRs/condition). The specific mean duration was chosen to enhance detection of task-related differences in the blood oxygen level-dependent (BOLD) signal ('activation') [39] and to allow sufficient time for sustained responses to become evident. Safety trials were similar, but terminated with the delivery of benign reinforcers (see below). Valence was continuously signaled during the anticipation epoch ('countdown') by the background color of the display. Temporal certainty was signaled by the nature of the integer stream. Certain trials always began with the presentation of the number 30. On uncertain trials, integers were randomly drawn from a near-uniform distribution ranging from 1 to 45 to reinforce the impression that they could be much shorter or longer than certain trials and to minimize incidental temporal learning ('time-keeping'). To concretely demonstrate the variable duration of uncertain trials, during scanning, the first three uncertain trials featured short (8.75 s), medium (15.00 s), and long (28.75 s) anticipation epochs. To mitigate potential confusion and eliminate mnemonic demands, a lower-case 'c' or 'u' was presented at the lower edge of the display throughout the anticipatory epoch. White-noise visual masks (3.2 s) were presented between trials to minimize the persistence of visual reinforcers in iconic memory.

Participants were periodically prompted (following the offset of the white-noise visual mask) to rate the intensity of fear/anxiety experienced a few seconds earlier, during the anticipation ('countdown') period of the prior trial, using a 1 (*minimal*) to 4 (*maximal*) scale and an MRI-compatible response pad (MRA, Washington, PA). Each condition was rated once per scan (16.7% trials). Premature ratings (<300 ms) were censored. All participants provided at least 6 usable ratings and rated each condition at least once. Skin conductance was continuously acquired throughout.

***Validation.*** Prior work by our group in the present sample and in an independent sample of community volunteers demonstrates that the threat conditions of the Maryland Threat Countdown task elicit robust symptoms of subjective distress and signs of objective arousal, confirming its validity as an experimental probe of fear and anxiety [5, 37] (**Supplementary Figure S2**).

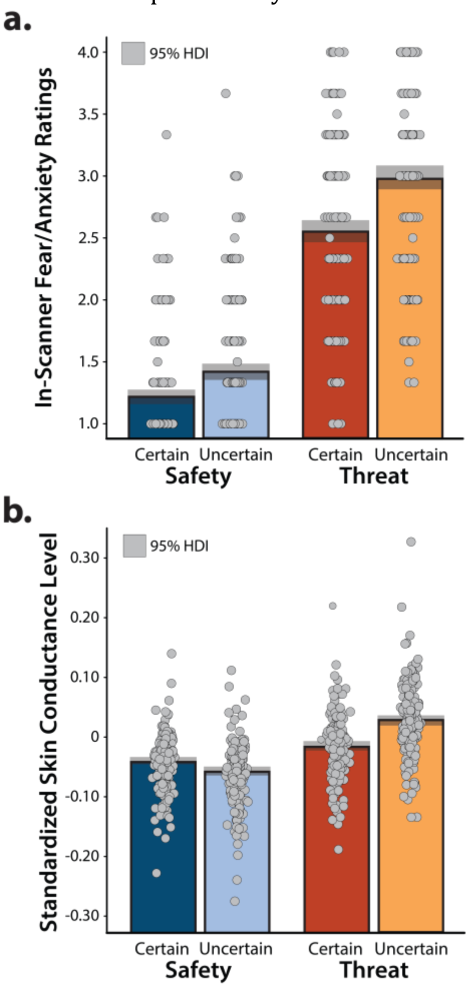

**Supplementary Figure S2. The Maryland Threat Countdown paradigm is a valid experimental probe of anticipatory fear and anxiety.** As detailed elsewhere, we used a series of repeated-measures GLMs to confirm that the threat-anticipation paradigm had the intended consequences for subjective distress and objective arousal [5]. **a.** ***Threat anticipation evokes subjective distress.*** Fearful and anxious feelings were significantly elevated during the anticipation of threat compared to safety, and this was particularly evident when threat encounters were uncertain in their timing (*Valence:* $F(1,219)=1{,}108.5$, $p<0.001$; *Certainty:* $F(1,219)=208.4$, $p<0.001$; *Valence × Certainty:* $F(1,219)=31.9$, $p<0.001$; *Threat, Uncertain vs. Certain:* $F(1,219)=148.5$, $p<0.001$; *Safety, Uncertain vs. Certain:* $F(1,219)=76.6$, $p<0.001$). **b.** ***Threat anticipation evokes objective signs of arousal.*** A similar pattern was evident for skin conductance level, an objective psychophysiological index of arousal (*Valence:* $F(1,217)=792.79$, $p<0.001$; *Certainty:* $F(1,217)=139.6$, $p<0.001$; *Valence × Certainty:* $F(1,217)=664.4$, $p<0.001$; *Threat, Uncertain vs. Certain:* $F(1,217)=457.8$, $p<0.001$; *Safety, Uncertain vs. Certain:* $F(1,217)=271.2$, $p<0.001$). Two participants were excluded from skin-conductance analyses due to insufficient usable data. Taken together, these observations confirm the validity of the threat-anticipation paradigm as an experimental probe of fear and anxiety, replicating prior work in community volunteers [37]. Bars indicate the means (*colored bars*), Bayesian 95% highest density intervals (HDI; *gray bands*), and individual participants (*gray dots*).

***Procedures.*** Prior to scanning, participants practiced an abbreviated version of the paradigm—without electrical stimulation—until they indicated and staff confirmed understanding. Benign and aversive electrical stimulation levels were individually titrated. *Benign stimulation.* Participants were asked whether they could "reliably detect" a 20 V stimulus and whether it was "at all unpleasant." If the subject could not detect the stimulus, the voltage was increased by 4 V and the process repeated. If the subject indicated that the stimulus was unpleasant, the voltage was reduced by 4V and the process was repeated. The final level chosen served as the benign electrical stimulation during the imaging assessment (*M*=21.06 V, *SD*=5.06). *Aversive stimulation.* Participants received a 100 V stimulus and were asked whether it was "as unpleasant as you are willing to tolerate"—an instruction specifically chosen to maximize anxious distress and arousal. If the subject indicated that they were willing to tolerate more intense stimulation, the voltage was increased by 10 V and the process repeated. If the subject indicated that the stimulus was too intense, the voltage was reduced by 5 V and the process repeated. The final level chosen served as the aversive electrical stimulation during the imaging assessment (*M*=117.85 V, *SD*=26.10). The intensity of aversive stimulation was weakly and negatively associated with individual differences in N/NE ($r(218)=-0.13$, $p=0.05$). Following each scan, staff re-assessed whether stimulation was sufficiently intense and increased the level as necessary.

***Electrical stimuli.*** Electrical stimuli (100 ms; 2 ms pulses every 10 ms) were generated using an MRI-compatible constant-voltage stimulator system (STMEPM-MRI; Biopac Systems, Inc., Goleta, CA). Stimuli were delivered using MRI-compatible, disposable carbon electrodes (Biopac) attached to the fourth and fifth digits of the non-dominant hand.

***Visual stimuli.*** A total of 72 aversive and benign photographs (1.8 s) were selected from the International Affective Picture System [12]. Visual stimuli were digitally back-projected (Powerlite Pro G5550, Epson America, Inc., Long Beach, CA) onto a semi-opaque screen mounted at the head-end of the scanner bore and viewed using a mirror mounted on the head-coil.

***Auditory stimuli.*** A total of 72 aversive and benign auditory stimuli (0.8 s) were adapted from open-access online sources. Auditory stimuli were delivered using an amplifier (PA-1 Whirlwind) with in-line noise-reducing filters and ear buds (S14; Sensimetrics, Gloucester, MA) fitted with noise-reducing ear plugs (Hearing Components, Inc., St. Paul, MN).

**Threat-related faces paradigm**

The threat-related faces paradigm takes the form of a pseudo-randomized block design and was administered in 2 scans, with a short break between scans (**Supplementary Figure S3**). During each scan, participants viewed standardized photographs of adults (half female) modeling prototypical angry faces, fearful faces, happy faces, or places (i.e., emotionally neutral everyday scenes; 7 blocks/condition/scan). To maximize signal strength and homogeneity and mitigate potential habituation [39-41], blocks consisted of 10 briefly presented photographs of faces or places (1.6 s) separated by fixation crosses (0.4 s). To further minimize potential habituation, each photograph was only presented a maximum of two times (for additional details, see [6]). To ensure engagement, participants judged whether the current photograph matched that presented on the prior trial (i.e., a '1-back' continuous performance task). Matches occurred 37.1% of the time.

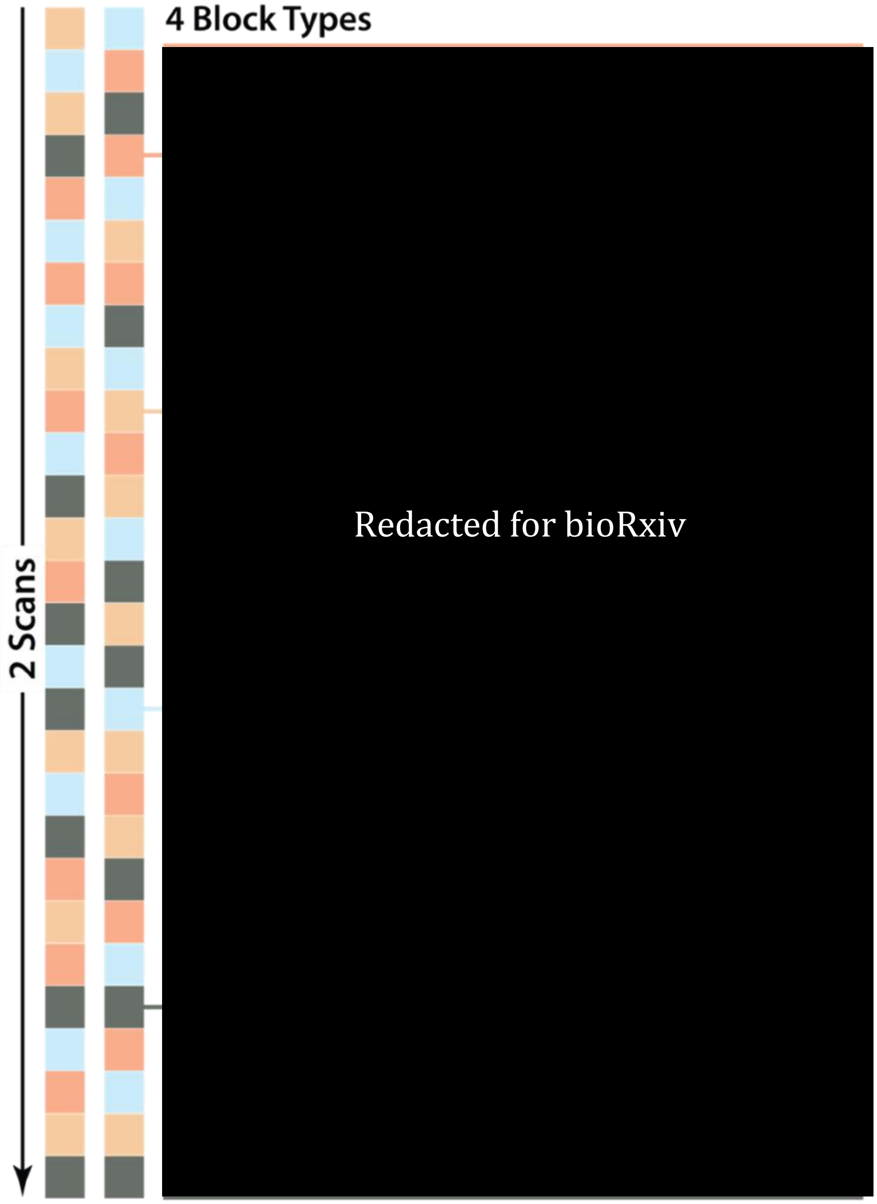

**Supplementary Figure S3. Emotional-faces paradigm.** The paradigm took the form of a pseudo-randomized block design and was administered in 2 scans, separated by a short break. During each scan, participants viewed standardized photographs of adults (half female) modeling prototypical angry faces, fearful faces, happy faces, or places (i.e., emotionally neutral everyday scenes; 7 blocks/condition/scan). Blocks consisted of 10 briefly presented photographs of faces or places (1.6 s) separated by fixation crosses (0.4 s). To further minimize potential habituation, each photograph was only presented a maximum of two times. To ensure engagement, participants judged whether the current photograph matched that presented on the prior trial (i.e., a '1-back' continuous performance task). Matches occurred 37.1% of the time.

#### MRI data acquisition

MRI data were acquired using a Siemens Magnetom TIM Trio 3 Tesla scanner (32-channel head-coil). During scanning, foam inserts were used to immobilize the participant's head within the head-coil and mitigate potential motion artifact. Participants were continuously monitored using an MRI-compatible eye-tracker (Eyelink 1000; SR Research, Ottawa, Ontario, Canada) and the AFNI real-time motion plugin [42]. Sagittal T1-weighted anatomical images were acquired using a magnetization prepared rapid acquisition gradient echo sequence (TR=2,400 ms; TE=2.01 ms; inversion time=1,060 ms; flip=8°; slice thickness=0.8 mm; in-plane=0.8 × 0.8 mm; matrix=300 × 320; field-of-view=240 × 256). A T2-weighted image was collected co-planar to the T1-weighted image (TR=3,200 ms; TE=564 ms; flip angle=120°). To enhance resolution, a multi-band sequence was used to collect oblique-axial EPI volumes (multiband acceleration=6; TR=1,250 ms; TE=39.4 ms; flip=36.4°; slice thickness=2.2 mm, number of slices=60; in-plane resolution=2.1875 × 2.1875 mm; matrix=96 × 96). Images were collected in the oblique-axial plane (approximately −20° relative to the AC-PC plane) to minimize potential susceptibility artifacts. For the threat-anticipation task, three 478-volume EPI scans were acquired. For the threat-perception (emotional-faces) task, two 454-volume EPI scans were acquired. The scanner automatically discarded 7 volumes prior to the first recorded volume. To enable fieldmap correction, two oblique-axial spin echo (SE) images were collected in opposing phase-encoding directions (rostral-to-caudal and caudal-to-rostral) at the same location and resolution as the functional volumes (i.e., co-planar; TR=7,220 ms; TE=73 ms). Measures of respiration and pulse were continuously acquired during scanning using a respiration belt and photo-plethysmograph affixed to the first digit of the non-dominant hand. Following the last scan, participants were removed from the scanner, debriefed, compensated, and discharged.

**MRI data processing pipeline**

Methods were optimized to minimize spatial normalization error and other potential sources of noise. Data were visually inspected before and after processing for quality assurance.

***Anatomical data processing.*** Methods are similar to those described in other recent reports by our group [5, 6, 12, 37]. T1-weighted images were inhomogeneity corrected using *N4* [43] and denoised using *ANTS* [44]. The brain was then extracted using *BEaST* [45] and brain-extracted and normalized reference brains from *IXI* [46]. Brain-extracted T1 images were normalized to a version of the brain-extracted 1-mm T1-weighted MNI152 (version 6) template [47] modified to remove extracerebral tissue. Normalization was performed using the diffeomorphic approach implemented in *SyN* (version 2.3.4) [44]. T2-weighted images were rigidly co-registered with the corresponding T1 prior to normalization. The brain extraction mask from the T1 was then applied. Tissue priors were unwarped to native space using the inverse of the diffeomorphic transformation [48]. Brain-extracted T1 and T2 images were segmented—using native-space priors generated in *FAST* (version 6.0.4) [49]—for subsequent use in T1-EPI co-registration (see below).

***Fieldmap data processing.*** SE images and *topup* were used to create fieldmaps. Fieldmaps were converted to radians, median-filtered, and smoothed (2-mm). The average of the distortion-corrected SE images was inhomogeneity corrected using *N4* and masked to remove extracerebral voxels using *3dSkullStrip* (version 19.1.00).

***Functional data processing.*** EPI files were de-spiked using *3dDespike*, slice-time corrected to the TR center using *3dTshift*, and motion corrected to the first volume and inhomogeneity corrected using *ANTS* (12-parameter affine). Transformations were saved in ITK-compatible format for subsequent use [50]. The first volume was extracted for EPI-T1 co-registration. The reference EPI volume was simultaneously co-registered with the corresponding T1-weighted image in native space and corrected for geometric distortions using boundary-based registration [49]. This step incorporated the previously created fieldmap, undistorted SE, T1, white matter (WM) image, and masks. The spatial transformations necessary to transform each EPI volume from native space to the reference EPI, from the reference EPI to the T1, and from the T1 to the template were concatenated and applied to the processed EPI data in a single step to minimize incidental spatial blurring. Normalized EPI data were resampled (2 $mm^3$) using fifth-order b-splines. To maximize anatomical resolution, no additional spatial filters were applied, consistent with prior work by our team and recent recommendations [5, 37, 51].

**fMRI data exclusions, modeling, and derivative variables**

***Data exclusions.*** Volume-to-volume displacement (>0.5 mm) was used to assess residual motion artifact. Scans with excessively frequent artifacts (>2 *SD*) were discarded. Participants with insufficient usable fMRI data (<2 scans of the threat-anticipation task or <1 scan of the threat-perception task) or who showed poor behavioral performance on the threat-perception task (accuracy <2 *SD*) were excluded from the relevant analyses (see above).

***Canonical first-level (single-subject) fMRI modeling.*** First- and second-level modeling procedures were identical to prior work by our group [5]. For each participant, first-level modeling was performed using general linear models (GLMs) implemented in *SPM12* (version 7771), with the default autoregressive model and the temporal band-pass filter set to the hemodynamic response function (HRF) and 128 s [52]. Regressors were convolved with a canonical HRF. *Threat-anticipation paradigm.* Hemodynamic reactivity was modeled using variable-duration rectangular ("boxcar") regressors that spanned the entirety of anticipation ('countdown') epochs of the uncertain threat, certain threat, and uncertain safety trials. To maximize design efficiency, certain-safety anticipation served as the reference condition and contributed to the baseline estimate [53]. Epochs corresponding to the presentation of the four types of reinforcers, white-noise visual masks, and rating prompts were simultaneously modeled using the same approach. EPI volumes acquired before the first trial and following the final trial were unmodeled and contributed to the baseline estimate. Nuisance variates included estimates of volume-to-volume displacement, motion (6 parameters × 3 lags), cerebrospinal fluid (CSF) signal, instantaneous pulse and respiration rates, and ICA-derived nuisance signals (e.g., brain edge, CSF edge, global motion, white matter) [54]. Volumes with excessive volume-to-volume displacement (>0.5 mm) and those during and immediately following reinforcer delivery were censored. *Threat-related faces paradigm.* Hemodynamic reactivity to blocks of each emotional expression (angry, fearful, and happy) was modeled using time-locked rectangular regressors. Place blocks served as the reference condition and contributed to the baseline estimate [53].

***Second-level (group) fMRI modeling.*** Standard whole-brain voxelwise ('second-level') repeated-measures GLMs ('random effects') were computed using *SPM12* and used to assess reactivity to the threat-anticipation and threat-related faces paradigms. For threat-anticipation, we modelled activation during the anticipation of threat (vs. safety), uncertain threat (vs. baseline), and certain threat (vs. baseline). For the threat-related faces, we modelled activation during the presentation of threat-related (angry and fearful) faces (vs. baseline). The decision to focus on estimates of activation (e.g., uncertain threat) relative to the implicit baseline (e.g., certain safety) was aimed at maximizing reliability [55-59].

***Validation.*** Prior work by our group using the present sample demonstrates that both tasks recruit the expected subcortical and cortical regions [5]. Key neuroimaging maps are publicly available at Neurovault (https://neurovault.org/collections/13109).

***Brain metrics.*** Consistent with prior work by our group, activation was quantified using spatially unsmoothed fMRI data, maximizing anatomical resolution [5, 37, 51]. *Threat-anticipation paradigm, subcortical regions.* Ce, BST, and PAG activation were quantified using established anatomical ROIs [51, 60, 61]. The BST ROI was modified to remove voxels encroaching upon neighboring regions of the striatum, thalamus, and ventricles. It mostly encompasses the supra-commissural BST, given the difficulty of reliably discriminating the borders of regions below the anterior commissure in T1-weighted images [62, 63]. Regression coefficients were averaged across voxels and hemispheres, separately for each combination of task contrast (e.g., uncertain threat), region (e.g., BST), and participant. Split-half parallel reliability ($\rho_{SP}$)—often termed Spearman-Brown corrected reliability—was acceptable (BST: 0.71-0.73; Ce: 0.67-0.71; PAG: 0.54-0.70) [64]. Similar values were evident for tau-equivalent reliability ($\rho_{ST}$). *Threat-anticipation paradigm, frontocortical regions.* Frontocortical regions are large and functionally heterogeneous, rendering anatomical ROIs suboptimal (e.g., MCC: 34,840 mm$^3$). To sidestep this issue, frontocortical ROIs were functionally prescribed based on peak task effects within anatomical regions identified in a large-scale neuroimaging study of threat anticipation [5], including MCC (Left: $x$=-8, $y$=16, $z$=34; Right: $x$=10, $y$=12, $z$=38), AI (Left: $x$=-34, $y$=12, $z$=8; Right: $x$=34, $y$=16, $z$=8), frontal operculum (FrO; Left: $x$=-32, $y$=20, $z$=10; Right: $x$=34, $y$=24, $z$=8), and dorsolateral prefrontal cortex/frontal pole (dlPFC/FP; Left: $x$=-30, $y$=52, $z$=28; Right: $x$=34, $y$=44, $z$=28). Anatomical regions were determined using the Harvard–Oxford atlas [65-67]. For each combination of region and hemisphere (e.g., left MCC), regression coefficients were extracted using cubical ROIs (216 $mm^3$) centered on the regional peak [6, 68]. To ensure invariant ROI prescriptions across conditions, peaks were identified using the voxelwise average of the three contrast maps (threat > safety, uncertain threat > certain safety, and certain threat > certain safety). Regression coefficients were separately averaged for each combination of condition, region, and participant. To minimize the number of comparisons, we created composite measures of frontocortical activity for each condition (i.e., averaged across regions; mean inter-region $r$=0.56; $\alpha$>0.79; $\rho_{SP}$=0.82-0.84). *Threat-related faces paradigm, amygdala.* The amygdala was anatomically defined using the Harvard-Oxford probabilistic atlas ($p$>0.25) [65-67]. Regression coefficients were extracted for each hemisphere for the threat-related faces contrast and averaged for each participant ($\rho_{SP}$=0.91).

#### Analytic strategy

***Overview.*** The overarching goal of the present study is to test the hypothesis that increased recruitment of canonical fear and anxiety circuitry during the anticipation of a genuinely distressing threat prospectively predicts a worsening trajectory of internalizing symptoms (IDAS Dysphoria) and the emergence or recurrence of internalizing disorders (SCID-5), that this association is more evident for uncertain than certain threat, and that these associations are amplified among individuals exposed to more severe or frequent NLEs (C-IRLE) during the longitudinal follow-up (**Figure 1** in the main report). We also explored whether these associations are evident for amygdala reactivity to threat-related faces.

In some research traditions and longitudinal analytic frameworks (e.g., cross-lagged panel models), 'trajectory' is reserved for contexts in which there are three or more timepoints available for analysis [69, 70]. Here, we use it in a broader sense to describe change from baseline to follow-up.

***Statistical software.*** Hypothesis testing was performed using standardized (*z*-transformed) predictors and models implemented in *R* (version 4.2.2) and *Rstudio* (version 2022.12.0.353) [71, 72]. Interaction terms were not standardized (though the predictor variables comprising the interaction were); thus coefficients reported are unstandardized. Data cleaning, reshaping, and analyses were performed using *car* (version 3.1.1) [73], *dplyr* (version 1.1.0) [74], *emmeans* (version 1.8.5) [75], *interactions* (version 1.1.5) [76], *MASS* (version 7.3.58.1) [77], *plyr* (version 1.8.8) [78], *ppcor* (version 1.1) [79], *psych* (version 2.3.6) [80], *readxl* (version 1.4.2) [81], *reshape2* (version 1.4.4) [82], *sandwich* (version 3.0.2) [83], *sensemakr* (version 0.1.4) [84], *stats* (version 4.2.2) [71], *tidyr* (version 1.3.0) [85], *tidyverse* (version 2.0.0) [86], *logspline* (version 2.1.19) [87], and *bayestestR* (version 0.13.0) [88]. Diagnostic procedures and data visualizations were used to confirm that test assumptions are satisfied [89]. Some figures were created using created using *gghalves* (version 0.1.4), *ggplot2* (version 3.4.1) [90], *raincloudplots* (version 0.2.0) [91], *MRIcron* (version 1.0.20190902) [92], and *MRIcroGL* (version 1.2.20201102) [92].

***Resource sharing.*** Statistical code and complete results for all analyses are available at OSF (https://osf.io/ufs7j/files/osfstorage).

***Hypothesis testing: broadband internalizing symptoms.*** Ordinary least-squares regression models were used to test whether increased recruitment of canonical threat circuitry is prospectively associated with the worsening of broadband internalizing symptoms and whether these associations are amplified among individuals exposed to more frequent or severe NLEs during the 30-month follow-up period. Each model controlled for baseline symptoms and took the following form:

(1) $Y_{Dysphoria\_IDAS\ Follow\text{-}Up} \sim b_0 + b_{Dysphoria\_IDAS\ Baseline} + b_{Brain\_Threat\_vs\_Safety} + b_{NLEs} + b_{NLEs*Brain\_Threat\_vs\_Safety} + \varepsilon$

Separate models were implemented for each brain metric (see above for details). The same approach was used to explore the predictive merits of amygdala reactivity to threat-related faces.

A second series of regression models was used to test whether these associations are more evident for temporally uncertain-threat anticipation. These models controlled for certain threat reactivity and took the following form:

(2) $Y_{Dysphoria\_IDAS\_Follow\text{-}Up} \sim b_0 + b_{Brain\_Uncertain\_Threat\_vs\_Certain\_Safety} + b_{Brain\_Certain\_Threat\_vs\_Certain\_Safety} + b_{Dysphoria\_IDAS\_Baseline} + b_{NLEs} + b_{NLEs*Brain\_Uncertain\_Threat\_vs\_Certain\_Safety} + b_{NLEs*Brain\_Certain\_Threat\_vs\_Certain\_Safety} + \varepsilon$

This provides an estimate of the variance in future internalizing symptoms uniquely explained by regional activation during the anticipation of temporally uncertain (and/or certain) threat. For all models, partial $R^2$ was reported for each predictor, allowing us to estimate how much residual variance in the outcome a predictor explained after partialing out the other variables in the model.

***Hypothesis testing: internalizing diagnoses.*** Logistic regression models were used to assess prospective associations with Any Internalizing Diagnosis. Because participants were free from prototypical internalizing illness at enrollment (see above for details), baseline diagnostic status was not included. These models took the following form:

(3) $Y_{Internalizing\_Diagnosis\_Follow\text{-}Up} \sim b_0 + b_{Brain\_Threat\_vs\_Safety} + b_{NLEs} + b_{NLEs*Brain\_Threat\_vs\_Safety} + \varepsilon$

(4) $Y_{Internalizing\_Diagnosis\_Follow\text{-}Up} \sim b_0 + b_{Brain\_Uncertain\_Threat\_vs\_Certain\_Safety} + b_{BST\_Certain\_Threat\_vs\_Certain\_Safety} + b_{NLEs} + b_{NLEs*Brain\_Uncertain\_Threat\_vs\_Certain\_Safety} + b_{NLEs*Brain\_Certain\_Threat\_vs\_Certain\_Safety} + \varepsilon$

As before, separate models were implemented for each brain metric and used to explore the predictive merits of amygdala reactivity to threat-related faces.

***Follow-up and exploratory analyses.*** To decompose significant interaction effects, simple slopes analyses were used [93]. This approach allowed us to disentangle how relations between brain activation and internalizing symptoms or diagnoses change at different levels of NLE exposure.

In direct comparisons of criterion validity, demographic variables, questionnaires, and other inexpensive traditional measures often outperform more sophisticated neuroimaging metrics [94]. To gauge the added explanatory value (i.e., incremental validity) of significant Brain-NLE interactions over measures of subjective distress and psychophysiological arousal, we recomputed the relevant regressions with an additional term for in-scanner ratings and skin conductance level (SCL), respectively. The specific contrasts used for the ratings and SCL variables were identical to those used for the relevant neuroimaging metric (e.g., threat vs. safety). As before, partial $R^2$ was reported for each model predictor.

Analyses revealed non-significant effects for the amygdala/threat-related faces task. Standardized Bayesian ($BF_{10}$) effect sizes were used to clarify interpretation [95]. $BF_{10}$ quantifies the relative performance of the null hypothesis ($H_0$; e.g., the absence of a credible mean difference) and the alternative hypothesis ($H_1$; e.g., the presence of a credible mean difference) on a 0 to ∞ scale. A key advantage of the Bayesian approach is that it can be used to formally quantify the relative strength of the evidence for $H_0$ ('test the null'), in contrast to standard null-hypothesis significance tests [96, 97]. It also does not require the data analyst to decide what constitutes a trivial difference, unlike traditional equivalence tests [98]. $BF_{10}$ was interpreted using established benchmarks [95]. Values <1 were interpreted as evidence of statistical equivalence (i.e., support for the null hypothesis), ranging from s*trong* ($BF_{10} \leq 0.10$), to *moderate* ($BF_{10}$=0.10-0.33), to *weak* ($BF_{10}$=0.33-1). The reciprocal of $BF_{10}$ represents the relative likelihood of the null hypothesis (e.g., $BF_{10}$=0.10, $H_0$ is 10 times more likely than $H_1$). Bayesian priors were set to a student's *t*-distribution with value $t$=0 to represent the null hypothesis, and this was contrasted with a posterior distribution with the *t* value from our model predictor. This method offers a standardized approach to quantifying evidence for/against the null based on prior and posterior samples of a single parameter [95].

### SUPPLEMENTARY RESULTS

#### Data missingness for questionnaire and interview assessments

**Supplementary Table S1.** Data missingness and exclusions for questionnaire and interview assessments.

| **Of the *n*=224 who completed one or both fMRI tasks:** | ***N*** | **%** |
|---|---|---|
| IDAS timepoints considered in isolation: | | |
| 0-Months: | 224 | 100.00% |
| 6-Months: | 222 | 99.11% |
| 24-Months: | 219 | 97.77% |
| 30-Months: | 218 | 97.32% |
| SCID-5 timepoints considered in isolation: | | |
| 0-Months: | 224 | 100.00% |
| 15-Months: | 221[a] | 98.66% |
| 30-Months: | 219 | 97.77% |
| C-IRLE timepoints considered in isolation: | | |
| 0-Months: | 224 | 100.00% |
| 15-Months: | 221[a] | 98.66% |
| 30-Months: | 218[b] | 97.32% |
| At least one Follow-Up IDAS [c]: | 220 | 98.21% |
| At least one Follow-Up SCID-5/C-IRLE [d]: | 222 | 99.11% |
| At least one Follow-Up for IDAS or SCID-5/C-IRLE: | 220 | 98.21% |

[a]One participant was excluded from present analyses due to incomplete 15-month SCID-5 and C-IRLE assessment data. [b]One participant was excluded from present analyses due to incomplete 30-month C-IRLE assessment data. [c]Only one Follow-Up assessment (24- and/or 30-month IDAS) was required for creation of the IDAS Dysphoria Follow-Up composite, and for inclusion in IDAS Dysphoria analyses. [d]Similarly, only one Follow-Up assessment (15- and/or 30-month SCID-5/C-IRLE) was required for inclusion in SCID-5/C-IRLE analyses. Abbreviations—C-IRLE, Cambridge Interview for Recent Life Events; IDAS, Inventory of Depression and Anxiety Symptoms; SCID-5, Structured Clinical Interview for *DSM-5*.

#### Regional reactivity to the fMRI paradigms

**Supplementary Table S2.** Regional reactivity to the threat-anticipation and threat-related faces paradigms.

| Task | Region | Contrast | *t* | *df* | *p* | Cohen's *d* |
|---|---|---|---|---|---|---|
| ***Threat-Anticipation*** | BST | Threat-Safety | 11.35 | 219 | <0.001 | 0.77 |
| | | Uncertain Threat[a] | 5.86 | 219 | <0.001 | 0.39 |
| | | Certain Threat[a] | 7.51 | 219 | <0.001 | 0.51 |
| | Ce | Threat-Safety | 8.10 | 219 | <0.001 | 0.55 |
| | | Uncertain Threat[a] | 3.08 | 219 | 0.002 | 0.21 |
| | | Certain Threat[a] | 4.39 | 219 | <0.001 | 0.30 |
| | PAG | Threat-Safety | 8.06 | 219 | <0.001 | 0.54 |
| | | Uncertain Threat[a] | 4.66 | 219 | <0.001 | 0.31 |
| | | Certain Threat[a] | 5.73 | 219 | <0.001 | 0.39 |
| | Frontocortical Composite | Threat-Safety | 20.47 | 219 | <0.001 | 1.38 |
| | | Uncertain Threat[a] | 17.33 | 219 | <0.001 | 1.17 |
| | | Certain Threat[a] | 9.98 | 219 | <0.001 | 0.67 |
| ***Threat-Related Faces*** | Amygdala | Faces-Places | 24.35 | 212 | <0.001 | 1.67 |

[a]Relative to the implicit baseline (i.e., Certain Safety anticipation). Abbreviations—BST, bed nucleus of the stria terminalis; Ce, dorsal amygdala in the region of the central nucleus; PAG, periaqueductal gray; ROI, region of interest. All *p*s ≤ 0.002.

#### Neural reactivity to the threat-versus-safety anticipation contrast was unrelated to regional changes in broadband internalizing symptoms

**Supplementary Table S3.** Linear regressions of follow-up Dysphoria on neural threat reactivity, NLEs, and their interaction, controlling for baseline Dysphoria.

| Model Terms [a,b] | *b* | *t* | *df* | *p* | $R^2_{partial}$ |
|---|---|---|---|---|---|
| ***Follow-Up Dysphoria*** | — | — | — | — | — |
| NLEs ** | 0.19 | 3.61 | 211 | 0.004 | 0.058 |
| Baseline Dysphoria *** | 0.63 | 12.00 | 211 | <0.001 | 0.406 |
| BST, Threat minus Safety Anticipation | -0.05 | -0.95 | 211 | 0.345 | 0.004 |
| BST TmS × NLEs | -0.01 | -0.18 | 211 | 0.857 | 0.000 |
| ***Follow-Up Dysphoria*** | — | — | — | — | — |
| NLEs *** | 0.20 | 3.93 | 211 | <0.001 | 0.068 |
| Baseline Dysphoria *** | 0.63 | 12.25 | 211 | <0.001 | 0.416 |
| Ce, Threat minus Safety Anticipation | -0.02 | -0.36 | 211 | 0.717 | 0.001 |
| Ce TmS × NLEs | 0.09 | 1.81 | 211 | 0.072 | 0.015 |
| ***Follow-Up Dysphoria*** | — | — | — | — | — |
| NLEs *** | 0.19 | 3.55 | 211 | <0.001 | 0.06 |
| Baseline Dysphoria *** | 0.62 | 11.95 | 211 | <0.001 | 0.404 |
| PAG, Threat minus Safety Anticipation | -0.01 | -0.27 | 211 | 0.788 | 0.000 |
| PAG TmS × NLEs | -0.07 | -1.17 | 211 | 0.244 | 0.006 |
| ***Follow-Up Dysphoria*** | — | — | — | — | — |
| NLEs *** | 0.19 | 3.72 | 211 | <0.001 | 0.062 |
| Baseline Dysphoria *** | 0.63 | 12.04 | 211 | <0.001 | 0.407 |
| Frontocortical Composite, TmS | -0.03 | -0.64 | 211 | 0.524 | 0.002 |
| Frontocortical Composite TmS × NLEs | 0.03 | 0.41 | 211 | 0.682 | 0.001 |

[a]Model intercept terms were not significant (*ps*>0.05). [b]All predictors were standardized (*z*-transformed). To sidestep standardizing the data twice, unstandardized coefficients are reported. Abbreviations—BST, bed nucleus of the stria terminalis; Ce, dorsal amygdala in the region of the central nucleus; PAG, periaqueductal gray; ROI, region of interest; TmS, Threat minus Safety Anticipation. * $p \leq 0.05$, ** $p<0.01$, *** $p<0.001$.

**BST and PAG reactivity to uncertain-threat demonstrate incremental validity over conventional measures of threat-elicited distress and psychophysiological arousal**

In head-to-head comparisons of criterion validity, demographic variables, questionnaires, and other simple measures often outperform more sophisticated and expensive neuroimaging metrics [94]. To gauge the added explanatory value (i.e., incremental validity) of significant Brain-NLE interactions over measures of subjective distress and psychophysiological arousal, we recomputed the relevant regressions with an additional term for in-scanner ratings and skin conductance level (SCL), respectively. To parallel the neuroimaging metrics, we computed the difference between uncertain-threat anticipation and the implicit baseline condition (certain safety) for in-scanner fear/anxiety ratings and skin conductance level. Using these derivative measures, results indicated that the $BST_{Uncertain\text{-}Threat}$-NLE interaction remained significant while controlling for either distress or arousal (*Controlling for Ratings:* $b$=-0.18, $t$(207)=-2.67, $p$=0.008; *Controlling for SCL:* $b$=-0.17, $t$(205)=-2.49, $p$=0.014). In contrast, the $PAG_{Uncertain\text{-}Threat}$-NLE interaction remained significant when controlling for distress, but not arousal (*Controlling for Ratings:* $b$=-0.14, $t$(207)=-2.08, $p$=0.039; *Controlling for SCL:* $b$=-0.13, $t$(205)=-1.84, $p$=0.068). Neither ratings nor arousal was associated with future internalizing symptoms, whether considered as main effects or as interactions with NLE exposure ($p$s>0.21). Taken together, these findings underscore the unique explanatory merits ("incremental validity") of the neural metrics in comparison to conventional measures of threat reactivity.

### Supplementary descriptive statistics for the models focused on broadband internalizing symptoms

**Supplementary Table S4.** Correlation matrix for variables used in the model predicting longitudinal changes in IDAS Dysphoria symptoms, based on BST reactivity to uncertain-/certain-threat anticipation and NLE exposure.

| Variable | *N* | *M* | *SD* | 1 | 2 | 3 | 4 | 5 |
|---|---|---|---|---|---|---|---|---|
| 1. NLEs | 216 | 10.83 | 4.37 | — | | | | |
| 2. Baseline Dysphoria | | 20.39 | 6.79 | 0.01 | — | | | |
| 3. Follow-Up Dysphoria | | 21.00 | 7.79 | 0.20** | 0.63*** | — | | |
| 4. BST, Uncertain-Threat Anticipation | | 0.33 | 0.85 | -0.07 | 0.14* | 0.07 | — | |
| 5. BST, Certain-Threat Anticipation | | 0.37 | 0.75 | 0.002 | 0.02 | -0.03 | 0.50*** | — |

Values reflect Pearson correlation coefficients. Abbreviations—BST, bed nucleus of the stria terminalis; NLEs, negative life events. * $p \leq 0.05$, ** $p < 0.01$, *** $p < 0.001$.

**Supplementary Table S5.** Partial correlation matrix for variables used in the model predicting longitudinal changes in IDAS Dysphoria symptoms, based on BST reactivity to uncertain-/certain-threat anticipation and NLE exposure, when controlling for baseline Dysphoria.

| Variable | *N* | 1 | 2 | 3 | 4 |
|---|---|---|---|---|---|
| 1. NLEs | 216 | — | | | |
| 2. Follow-Up Dysphoria | | 0.25*** | — | | |
| 3. BST, Uncertain-Threat Anticipation | | -0.08 | -0.02 | — | |
| 4. BST, Certain-Threat Anticipation | | 0.002 | -0.06 | 0.50*** | — |

Values reflect Pearson partial correlation coefficients ***when controlling for Baseline Dysphoria.*** Abbreviations—BST, bed nucleus of the stria terminalis; NLEs, negative life events. * $p \leq 0.05$, ** $p < 0.01$, *** $p < 0.001$.

**Supplementary Table S6.** Correlation matrix for variables used in the model predicting longitudinal changes in IDAS Dysphoria symptoms, based on PAG reactivity to uncertain-/certain-threat anticipation and NLE exposure.

| Variable | *N* | *M* | *SD* | 1 | 2 | 3 | 4 | 5 |
|---|---|---|---|---|---|---|---|---|
| 1. NLEs | 216 | 10.83 | 4.37 | — | | | | |
| 2. Baseline Dysphoria | | 20.39 | 6.79 | 0.01 | — | | | |
| 3. Follow-Up Dysphoria | | 21.00 | 7.79 | 0.20** | 0.63*** | — | | |
| 4. PAG, Uncertain-Threat Anticipation | | 0.38 | 1.23 | -0.09 | -0.15* | -0.06 | — | |
| 5. PAG, Certain-Threat Anticipation | | 0.47 | 1.26 | -0.08 | -0.14* | -0.14* | 0.59*** | — |

Values reflect Pearson correlation coefficients. Abbreviations—NLEs, negative life events; PAG, periaqueductal gray. * $p \leq 0.05$, ** $p < 0.01$, *** $p < 0.001$.

**Supplementary Table S7.** Partial correlation matrix for variables used in the model predicting longitudinal changes in IDAS Dysphoria symptoms, based on PAG reactivity to uncertain-/certain-threat anticipation and NLE exposure, when controlling for baseline Dysphoria.

| Variable | *N* | 1 | 2 | 3 | 4 |
|---|---|---|---|---|---|
| 1. NLEs | 216 | — | | | |
| 2. Follow-Up Dysphoria | | 0.25*** | — | | |
| 3. PAG, Uncertain-Threat Anticipation | | -0.09 | 0.04 | — | |
| 4. PAG, Certain-Threat Anticipation | | -0.08 | -0.07 | 0.58*** | — |

Values reflect Pearson partial correlation coefficients ***when controlling for Baseline Dysphoria.*** Abbreviations—NLEs, negative life events; PAG, periaqueductal gray. * *p*≤0.05, ** *p*<0.01, *** *p*<0.001.

**Supplementary Table S8.** Correlation matrix for variables used in the model predicting longitudinal changes in IDAS Dysphoria symptoms, based on Ce reactivity to uncertain-/certain-threat anticipation and NLE exposure.

| Variable | *N* | *M* | *SD* | 1 | 2 | 3 | 4 | 5 |
|---|---|---|---|---|---|---|---|---|
| 1. NLEs | 216 | 10.83 | 4.37 | — | | | | |
| 2. Baseline Dysphoria | | 20.39 | 6.79 | 0.01 | — | | | |
| 3. Follow-Up Dysphoria | | 21.00 | 7.79 | 0.20** | 0.63*** | — | | |
| 4. Ce, Uncertain-Threat Anticipation | | 0.10 | 0.53 | -0.02 | -0.04 | -0.05 | — | |
| 5. Ce, Certain-Threat Anticipation | | 0.14 | 0.48 | 0.001 | -0.05 | -0.09 | 0.65*** | — |

Values reflect Pearson correlation coefficients. Abbreviations—NLEs, negative life events; Ce, central nucleus of the amygdala. * *p*≤0.05, ** *p*<0.01, *** *p*<0.001.

**Supplementary Table S9.** Partial correlation matrix for variables used in the model predicting longitudinal changes in IDAS Dysphoria symptoms, based on Ce reactivity to uncertain-/certain-threat anticipation and NLE exposure, when controlling for baseline Dysphoria.

| Variable | *N* | 1 | 2 | 3 | 4 |
|---|---|---|---|---|---|
| 1. NLEs | 216 | — | | | |
| 2. Follow-Up Dysphoria | | 0.25*** | — | | |
| 3. Ce, Uncertain-Threat Anticipation | | -0.02 | -0.03 | — | |
| 4. Ce, Certain-Threat Anticipation | | 0.001 | -0.07 | 0.65*** | — |

Values reflect Pearson partial correlation coefficients ***when controlling for Baseline Dysphoria.*** Abbreviations—NLEs, negative life events; Ce, centra l nucleus of the amygdala. * *p*≤0.05, ** *p*<0.01, *** *p*<0.001.

**Supplementary Table S10.** Correlation matrix for variables used in the model predicting longitudinal changes in IDAS Dysphoria symptoms, based on frontocortical composite reactivity to uncertain-/certain-threat anticipation and NLE exposure.

| Variable | *N* | *M* | *SD* | 1 | 2 | 3 | 4 | 5 |
|---|---|---|---|---|---|---|---|---|
| 1. NLEs | 216 | 10.83 | 4.37 | — | | | | |
| 2. Baseline Dysphoria | | 20.39 | 6.79 | 0.01 | — | | | |
| 3. Follow-Up Dysphoria | | 21.00 | 7.79 | 0.20** | 0.63*** | — | | |
| 4. Frontocortical Composite, Uncertain-Threat Anticipation | | 0.94 | 0.80 | -0.16* | 0.17* | 0.09 | — | |
| 5. Frontocortical Composite, Certain-Threat Anticipation | | 0.45 | 0.66 | -0.07 | 0.07 | -0.01 | 0.48*** | — |

Values reflect Pearson correlation coefficients. Abbreviations—NLEs, negative life events. * *p*≤0.05, ** *p*<0.01, *** *p*<0.001.

**Supplementary Table S11.** Partial correlation matrix for variables used in the model predicting longitudinal changes in IDAS Dysphoria symptoms, based on frontocortical composite to uncertain-/certain-threat anticipation and NLE exposure, when controlling for baseline Dysphoria.

| Variable | *N* | 1 | 2 | 3 | 4 |
|---|---|---|---|---|---|
| 1. NLEs | 216 | — | | | |
| 2. Follow-Up Dysphoria | | 0.25*** | — | | |
| 3. Frontocortical Composite, Uncertain-Threat Anticipation | | -0.17* | -0.02 | — | |

| | | | | | |
|---|---|---|---|---|---|
| 4. Frontocortical Composite, Certain-Threat Anticipation | | -0.07 | -0.06 | 0.48*** | — |

Values reflect Pearson partial correlation coefficients ***when controlling for Baseline Dysphoria.*** Abbreviations—NLEs, negative life events. * *p*≤0.05, ** *p*<0.01, *** *p*<0.001.

**Supplementary Table S12.** Correlation matrix for variables used in the model predicting longitudinal changes in IDAS Dysphoria symptoms, based on amygdala reactivity to threat-related faces and NLE exposure.

| **Variable** | ***N*** | ***M*** | ***SD*** | 1 | 2 | 3 | 4 |
|---|---|---|---|---|---|---|---|
| 1. NLEs | 209 | 10.81 | 4.36 | — | | | |
| 2. Baseline Dysphoria | | 20.32 | 6.73 | 0.04 | — | | |
| 3. Follow-Up Dysphoria | | 20.99 | 7.68 | 0.21** | 0.63*** | — | |
| 4. Amygdala, Faces minus Places | | 0.69 | 0.42 | -0.11 | 0.003 | -0.08 | — |

Values reflect Pearson correlation coefficients. * *p*≤0.05, ** *p*<0.01, *** *p*<0.001.

**Supplementary Table S13.** Partial correlation matrix for variables used in the model predicting longitudinal changes in IDAS Dysphoria symptoms, based on amygdala reactivity to threat-related faces and NLE exposure, when controlling for baseline Dysphoria.

| **Variable** | ***N*** | 1 | 2 | 3 |
|---|---|---|---|---|
| 1. NLEs | 209 | — | | |
| 2. Follow-Up Dysphoria | | 0.24*** | — | |
| 3. Amygdala, Faces minus Places | | -0.11 | -0.10 | — |

Values reflect Pearson partial correlation coefficients ***when controlling for Baseline Dysphoria.*** Abbreviations—NLEs, negative life events. * *p*≤0.05, ** *p*<0.01, *** *p*<0.001.

#### Increased NLE exposure confers risk for future internalizing diagnoses

Paralleling the approach we used for dimensional internalizing symptoms, we implemented a simplified logistic regression model (or “base model”) to confirm that NLE exposure is associated with increased odds of being diagnosed with a new or recurrent illness during the 30-month follow-up ($b$=0.97, $z$=4.83, $p$<0.001, odds ratio (OR)=2.65, 95% CI[1.78, 3.93]).

#### Individual differences in neural reactivity are unrelated to future internalizing diagnoses

Variation in regional reactivity to the two fMRI tasks was not significantly related to the emergence of a clinically diagnosed internalizing disorder over the follow-up period, regardless of NLE exposure ($p$s>0.07; **Supplementary Tables 14-16**).

**Supplementary Table S14.** Logistic regressions of a new or recurrent internalizing diagnosis during follow-up on neural threat reactivity, NLEs, and their interaction.

| Model Terms [a,b] | *b* | *z* | *p* | *OR* | *OR 95% CI* | *Probability* |
|---|---|---|---|---|---|---|
| ***New or Recurrent Internalizing Dx*** | — | — | — | — | — | — |
| NLEs *** | 0.98 | 4.85 | <0.001 | 2.68 | [1.80, 3.99] | 72.81% |
| BST, Threat minus Safety Anticipation | -0.11 | -0.58 | 0.565 | 0.89 | [0.61, 1.31] | 47.19% |
| BST TmS × NLEs | -0.03 | -0.16 | 0.872 | 0.97 | [0.69, 1.37] | 49.30% |
| ***New or Recurrent Internalizing Dx*** | — | — | — | — | — | — |
| NLEs *** | 1.01 | 4.81 | <0.001 | 2.74 | [1.82, 4.13] | 73.26% |
| Ce, Threat minus Safety Anticipation | 0.00 | 0.02 | 0.988 | 1.00 | [0.70, 1.43] | 50.07% |
| Ce TmS × NLEs | -0.35 | -1.76 | 0.079 | 0.71 | [0.48, 1.04] | 41.38% |
| ***New or Recurrent Internalizing Dx*** | — | — | — | — | — | — |
| NLEs *** | 0.94 | 4.63 | <0.001 | 2.56 | [1.72, 3.81] | 71.89% |
| PAG, Threat minus Safety Anticipation | -0.27 | -1.38 | 0.167 | 0.77 | [0.52, 1.12] | 43.36% |
| PAG TmS × NLEs | -0.08 | -0.38 | 0.701 | 0.92 | [0.60, 1.40] | 47.94% |
| ***New or Recurrent Internalizing Dx*** | — | — | — | — | — | — |
| NLEs *** | 1.02 | 4.84 | <0.001 | 2.77 | [1.83, 4.18] | 73.46% |
| Frontocortical Composite, TmS | 0.22 | 1.24 | 0.215 | 1.25 | [0.88, 1.77] | 55.49% |
| Frontocortical Composite TmS x NLEs | -0.26 | -1.40 | 0.162 | 0.77 | [0.54, 1.11] | 43.62% |

[a]Model intercept terms not reported were all statistically significant (*p*s<0.001). [b]All predictors were standardized (*z*-transformed). To sidestep standardizing the data twice, unstandardized coefficients are reported. Abbreviations—BST, bed nucleus of the stria terminalis; Ce, dorsal amygdala in the region of the central nucleus; CI, confidence interval; Dx, diagnosis; OR, odds ratio; PAG, periaqueductal gray; TmS, Threat minus Safety anticipation. * *p*≤0.05, ** *p*<0.01, *** *p*<0.001.

**Supplementary Table S15.** Logistic regressions of a new or recurrent internalizing disorder during follow-up on certain and uncertain threat reactivity, NLEs, and their interactions.

| Model Terms [a,b] | *b* | *z* | *p* | *OR* | *OR 95% CI* | *Probability* |
|---|---|---|---|---|---|---|
| ***New or Recurrent Internalizing Dx*** | — | — | — | — | — | — |
| NLEs *** | 0.98 | 4.74 | <0.001 | 2.66 | [1.78, 4.00] | 72.70% |
| BST, Uncertain-Threat Anticipation[c] | -0.09 | -0.39 | 0.695 | 0.92 | [0.59, 1.42] | 47.82% |
| BST, Certain-Threat Anticipation[c] | -0.07 | -0.34 | 0.735 | 0.93 | [0.61, 1.42] | 48.16% |
| BST UT[c] × NLEs | 0.09 | 0.40 | 0.689 | 1.09 | [0.71, 1.68] | 52.21% |
| BST CT[c] × NLEs | -0.13 | -0.56 | 0.578 | 0.87 | [0.55, 1.40] | 46.65% |
| ***New or Recurrent Internalizing Dx*** | — | — | — | — | — | — |
| NLEs *** | 1.02 | 4.74 | <0.001 | 2.76 | [1.82, 4.21] | 73.44% |
| Ce, Uncertain-Threat Anticipation[c] | -0.24 | -0.92 | 0.357 | 0.79 | [0.47, 1.31] | 43.99% |
| Ce, Certain-Threat Anticipation[c] | 0.19 | 0.74 | 0.458 | 1.21 | [0.73, 2.00] | 54.74% |
| Ce UT[c] × NLEs | -0.09 | -0.32 | 0.752 | 0.91 | [0.52, 1.60] | 47.74% |
| Ce CT[c] × NLE | -0.28 | -0.96 | 0.340 | 0.76 | [0.43, 1.34] | 43.15% |
| ***New or Recurrent Internalizing Dx*** | — | — | — | — | — | — |
| NLEs *** | 0.95 | 4.57 | <0.001 | 2.58 | [1.72, 3.87] | 72.05% |
| PAG, Uncertain-Threat Anticipation[c] | -0.32 | -1.38 | 0.166 | 0.72 | [0.46, 1.14] | 41.98% |
| PAG, Certain-Threat Anticipation[c] | 0.10 | 0.43 | 0.670 | 1.11 | [0.70, 1.75] | 52.51% |
| PAG UT[c] × NLEs | -0.02 | -0.09 | 0.930 | 0.98 | [0.60, 1.61] | 49.44% |
| PAG CT[c] × NLEs | -0.07 | -0.29 | 0.771 | 0.93 | [0.59, 1.48] | 48.30% |
| ***New or Recurrent Internalizing Dx*** | — | — | — | — | — | — |
| NLEs *** | 1.03 | 4.90 | <0.001 | 2.79 | [1.85, 4.21] | 73.62% |
| Frontocortical Composite, UT[c] | 0.03 | 0.15 | 0.881 | 1.03 | [0.68, 1.56] | 50.79% |
| Frontocortical Composite, CT[c] | 0.26 | 1.28 | 0.200 | 1.29 | [0.87, 1.91] | 56.36% |
| Frontocortical Composite UT[c] × NLEs | -0.13 | -0.59 | 0.557 | 0.88 | [0.57, 1.35] | 46.78% |
| Frontocortical Composite CT[c] × NLEs | 0.08 | 0.35 | 0.730 | 1.08 | [0.70, 1.65] | 51.88% |

[a]Model intercept terms not reported were all statistically significant (*p*s<0.001). [b]All predictors were standardized (*z*-transformed). To sidestep standardizing the data twice, unstandardized coefficients are reported. [c]Relative to the implicit baseline (i.e., Certain-Safety anticipation). Abbreviation—NLEs, negative life events, Abbreviations—BST, bed nucleus of the stria terminalis; Ce, dorsal amygdala in the region of the central nucleus; CI, confidence interval; CT, Certain-Threat anticipation; OR, odds ratio; PAG, periaqueductal gray; UT, Uncertain-Threat anticipation. * *p*≤0.05, ** *p*<0.01, *** *p*<0.001.

**Supplementary Table S16.** Logistic regression of a new or recurrent internalizing disorder during follow-up on amygdala reactivity to threat-related faces, NLEs, and their interaction.

| Model Terms [a,b] | *b* | *z* | *p* | *OR* | *OR 95% CI* | *Probability* |
|---|---|---|---|---|---|---|
| ***New or Recurrent Internalizing Dx*** | — | — | — | — | — | — |
| NLEs *** | 0.94 | 4.47 | <0.001 | 2.56 | [1.70, 3.87] | 71.92% |
| Amygdala, Faces minus Places | -0.11 | -0.57 | 0.568 | 0.90 | [0.61, 1.31] | 47.24% |
| Amygdala, Faces minus Places × NLEs | -0.27 | -1.26 | 0.209 | 0.77 | [0.51, 1.16] | 43.40% |

[a]Model intercept term was statistically significant ($p<0.001$). [b]All predictors were standardized ($z$-transformed). To sidestep standardizing the data twice, unstandardized coefficients are reported. Abbreviations—CI, confidence interval; OR, odds ratio. * $p\leq0.05$, ** $p<0.01$, *** $p<0.001$.

#### SUPPLEMENTARY DISCUSSION

##### Magnitude of prospective-longitudinal associations

$BST/PAG_{Uncertain\text{-}Threat}$-NLE associations explain 2.1-3.4% of the variance in future internalizing symptoms (above-and-beyond variance explained by baseline symptoms). The magnitude of these two prospective associations—while far too modest for clinical or treatment-development purposes—appears plausible, given the complexity of BST/PAG functional neuroanatomy and the multitude of factors influencing both self-reported internalizing symptoms and the BOLD signal [99-103]. It compares favorably with other psychiatrically relevant neurobiological associations, including prospective associations between ventral-striatum reward-reactivity and depression (1%) [104]. From a mechanistic perspective, the small-but-reliable "hits" uncovered by adequately powered association studies are useful for prioritizing targets in animal models and human neuromodulation studies. This reflects the fact that modest brain-behavior associations do not preclude much larger effects with targeted biological interventions [94, 105].

##### Absence of associations with *DSM-5* diagnoses

The present results underscore the utility of transdiagnostic, dimensional approaches to internalizing illness. They are consistent with meta-analytic evidence that prospective associations are consistently weaker for categorical diagnoses than dimensional measures of internalizing illness [27, 106-108]. Our findings do not support clear inferences about the cause of this discrepancy, which might reflect the greater heterogeneity of diagnostic categories, the loss of information due to arbitrary clinical boundaries, or reduced psychometric reliability [27].

##### Detailed Limitations

Clearly, several key challenges remain for the future. First, it will be important to determine whether our conclusions generalize to more demographically representative samples, other types of experimental threat (e.g., social), and other kinds of threat uncertainty (e.g., probability, risk, ambiguity). It merits comment that the absence of reward trials precludes strong claims about valence. While unlikely, similar associations might be evident for uncertain-reward anticipation. It will also be useful to explore prospective associations with narrower symptoms of internalizing illness, including anxious arousal [10, 109, 110]. From the perspective of understanding the development of PTSD, it will be fruitful to examine samples exposed to severe NLEs [e.g., 111]. Second, the absence of significant associations with *DSM-5* diagnoses precludes strong inferences about impairment. Moving forward, it will be important to determine whether variation in BST/PAG function are associated with measures of daily function. It might also be informative to examine the importance of unmodeled nuisance variates (e.g., longitudinal changes in treatment status). Third, our sample was powered to detect effects as small as the BST-NLEs interaction. While statistically significant, the PAG-NLEs interaction did not reach this benchmark, underscoring the need for interpretative caution and empirical replication. Fourth, although our interview-based NLE assessment dovetails with methodological recommendations [28], future work may benefit from more intensive sampling (which would enhance temporal resolution and minimize recall errors and biases); from disaggregating stressor type, frequency, and severity; from clarifying temporal precedence and independence from internalizing illness; and from blinding interviewers to psychiatric history. Fifth, the effects of stress, adversity, and trauma are multidimensional, nonlinear, interactive, and age-dependent [112]. Unraveling this complexity is an important avenue for future research, but it will require well-powered longitudinal cohorts and more comprehensive assessments. Sixth, the BST and PAG are complex and can be subdivided into multiple subdivisions, each containing intermingled cell types with distinct, even opposing functions [113-115]. Animal models will be critical for generating testable hypotheses about the most relevant molecules, cell types, and microcircuits. Seventh, the etiology of internalizing psychopathology is complex, multifactorial, and likely reflects the coordinated interactions of widely distributed neural networks [116-119]. Moving forward, it will be important to clarify the relevance of functional connectivity between the BST, PAG, and other regions, and assess the predictive merits of multivoxel composites ("signatures") of anxiety-related brain activity [11, 120-122]. Multivoxel signatures provide a principled means of assessing the overall reactivity of functionally defined, psychiatrically relevant neural circuits [123] and often provide greater reliability and power than conventional voxelwise or ROI approaches [124, 125].
